# Airway microbiome–host transcriptome networks link microbial dysbiosis to survival outcomes and therapeutic opportunities in severe COVID-19

**DOI:** 10.1101/2025.06.24.661228

**Authors:** Kai-Pu Chen, Chen-Hao Huang, Kuan-Ya Chen, Chia-Lang Hsu, Hsuan-Cheng Huang, Hsueh-Fen Juan

## Abstract

The lower respiratory tract microbiome may critically influence outcomes in severe COVID-19, yet remains understudied. We analyzed bronchoalveolar lavage fluid from 86 severe patients requiring mechanical ventilation, stratified into two groups. We identified 213 bacteria with differential abundance, 11 bacteria associated with differentially expressed genes, and 39 of them were associated with patient survival. The more serious group exhibited reduced microbial network complexity, suggesting ecosystem disruption. To explore therapeutic potential, we employed our previously published model and predicted that compounds like palbociclib, XMD-1499, RS-17053, and RS-504393 may shift host gene expression favoring beneficial microbial signatures. These results highlight an important link between microbial dysbiosis and COVID-19 severity, offering promising directions for microbiome-informed therapeutic strategies. Our findings underscore the importance of the lung microbiome in modulating host responses and clinical outcomes, and suggest that modulating microbial-host interactions may serve as a novel adjunctive approach in the treatment of severe COVID-19.

**Graphical abstract:** 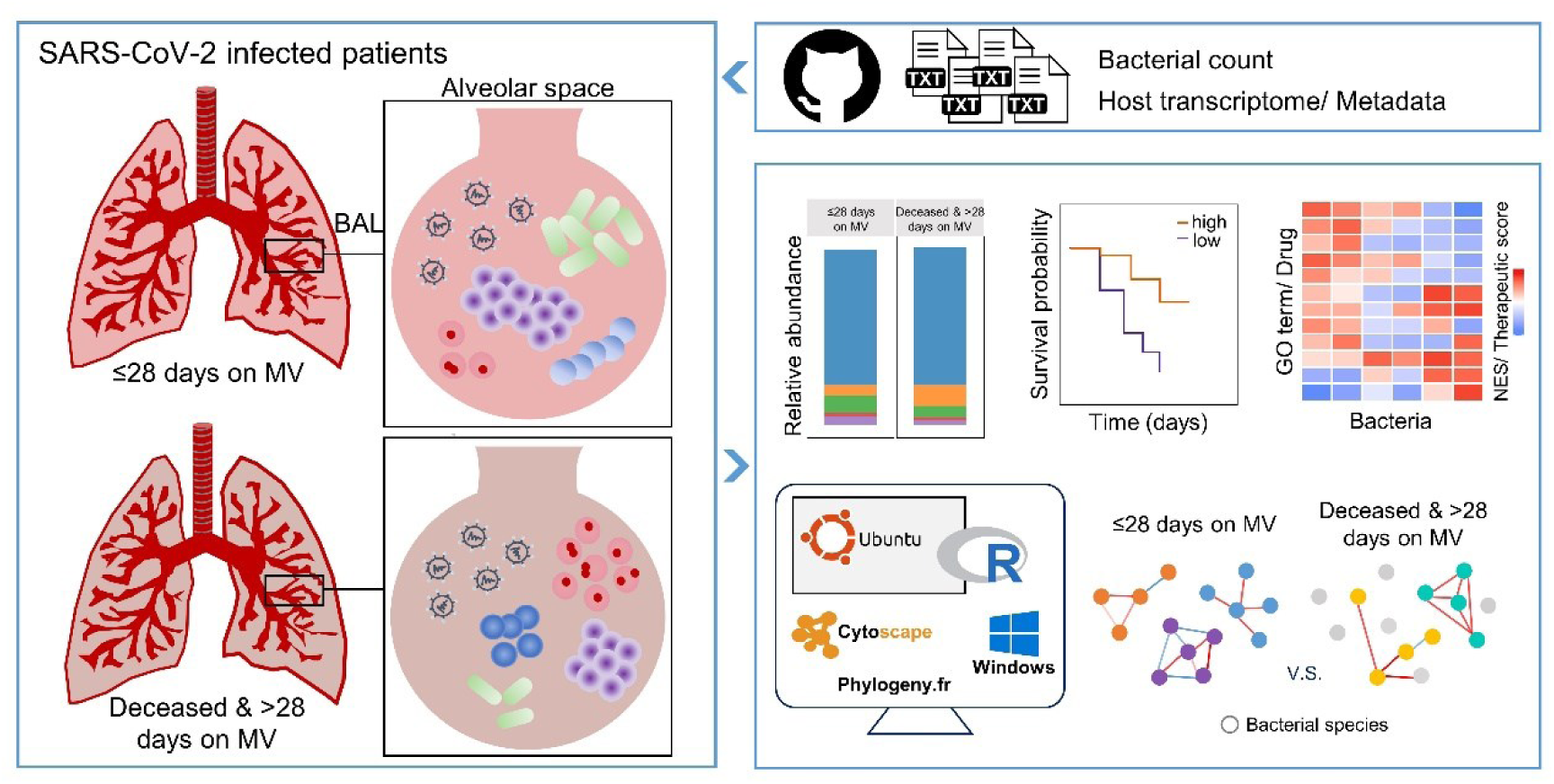

## INTRODUCTION

Since the outbreak of the COVID-19 pandemic in 2019 until June 9, 2024, resulting from the infection of severe acute respiratory syndrome coronavirus 2 (SARS-CoV-2), World Health Organization (WHO) has documented more than 775 million cases of COVID-19 and 7.05 million deaths worldwide.^1^ Most infected patients experience mild symptoms, such as cough, fever, fatigue, or myalgia. There are also individuals who are asymptomatic and can still transmit the virus, which complicates efforts to control the pandemic.^2^ SARS-CoV-2 can also cause severe and life-threatening conditions, including interstitial pneumonia, respiratory failure, acute respiratory distress syndrome (ARDS), and even multiorgan failure.^3,4^ Previous studies have identified various factors that contribute to the severity of COVID-19 patients. These factors include age, obesity, and immune response, such as dysregulated levels of cytokines and chemokines.^5,6^ Moreover, studies have shown that microbes present in oral and respiratory tract can impact specific immune cells, potentially related to the usage of mechanical ventilation.^7,8^

Previous studies have detected pathogenic bacteria in individuals with severe pneumonia, including those with ventilator-associated pneumonia (VAP).^9–12^ The primary site of disease activity caused by SARS-CoV-2 is the alveolar space within the lower respiratory tract. However, limited attention has been paid to investigating the microbiome of the lower respiratory site in severe COVID-19 patients with ventilator support. Previously, bronchoalveolar lavage (BAL) analysis has been used to characterize pneumonia caused by viruses and bacteria, identify the presence of a secondary infection, or ascertain if a non-infectious factor is contributing to the progression of severe pulmonary illness.^13–15^ Sulaiman et al. conducted a study analyzing the BAL derived microbiome in the lower respiratory tract of patients having mechanical ventilation; the findings include that a fatal outcome was associated with an enriched *Mycoplasma salivarium*, high SARS-CoV-2 load, low anti-SARS-CoV-2 antibody levels, among other factors.^16^

Patients were originally categorized into three distinct groups based on their duration on mechanical ventilation (MV): those on MV for ≤28 days (the ≤28 days on MV group), those on MV for >28 days (the >28 days on MV group), and those who unfortunately passed away during the study (the deceased group). We found that there were few host differentially expressed genes (DEGs) between groups of >28 days on MV and deceased. Additionally, we observed low gene expression in the host transcriptomic data of certain patients. Consequently, we excluded these samples from our analysis and merged the patients from the >28 days on MV group with those from the deceased group, forming the deceased & >28 days on MV group. We aimed to identify the specific bacterial species associated with dysbiosis in critically ill COVID-19 patients within the two distinct groups, and discover bacteria that impact patient survival as well as potential medications for treatment. We first investigated bacterial composition, differential abundance of bacterial species, diversity indices and bacterial co-abundance networks in two simplified clinical groups. Spearman correlation coefficient (SCC) was used to calculate the correlations between bacterial relative abundance and host gene expression. Additionally, we used Fisher’s exact test to identify nine plus two potentially DEGs-associated bacterial species. Log-rank test was employed to determine survival-related bacterial species from the nine plus two potentially DEGs-associated or 213 differentially abundant bacterial species. The biological functions of these important bacteria were inferred from FGSEA using the input that maintained the original rank in descending order of SCC and randomized once and sorted SCC values of every interesting bacterium. We calculated Sparse Correlations for Compositional data (SparCC) in the bacterial co-abundance networks as it provides the best linear relationship inference and better precision with highly compositional data (Inverse Simpson n_eff_ < 13), as shown by Weiss et al.^17^ Molecular Complex Detection (MCODE), a Cytoscape plugin, was used to discover modules with highly connected local regions, and the networks with highlighted nodes and edges were displayed using Cytoscape.^18,19^ Finally, our gene-perturbation-based drug prediction identified compounds targeting DEGs correlated with the survival-related bacterial species.

## RESULTS

### Overview of the analytical workflow

The analytical workflow of this study is depicted in Figure 1. To explore the impact of bacterial species on critically ill COVID-19 patients, we utilized data files containing bacterial count, metadata and host transcriptome matrices from the segalmicrobiomelab’s GitHub repository.^16^ This study cohort comprised 86 severely ill COVID-19 patients, divided into two clinical groups based on the host transcriptome: “≤ 28 days on MV” (*N* = 21) and “Deceased & >28 days on MV” (*N* = 65). An overview of patient information is presented in Table 1, with additional details available in Table S1. Given the DEGs identified in the originally published work indicated minimal differential expression between patients who had either died or received MV for more than 28 days, these patients were consolidated into a single group labeled “Deceased & >28 days on MV”. The refined and normalized host transcriptomic matrix included 86 BAL samples and 16282 genes (Figure S1 and Table S2). The microbial profiles comprised 4157 species, 1261 genera and 38 phyla within the 86 matched BAL samples. We conducted a comparative analysis of bacterial composition, diversity, bacterial relative abundance (using CLR-transformation: see Table S3), bacterial co-abundance and host gene expression between two clinical outcome groups, aiming to identify the role of bacterial species and host DEGs in the critically ill COVID-19 patients.

**Figure 1.**
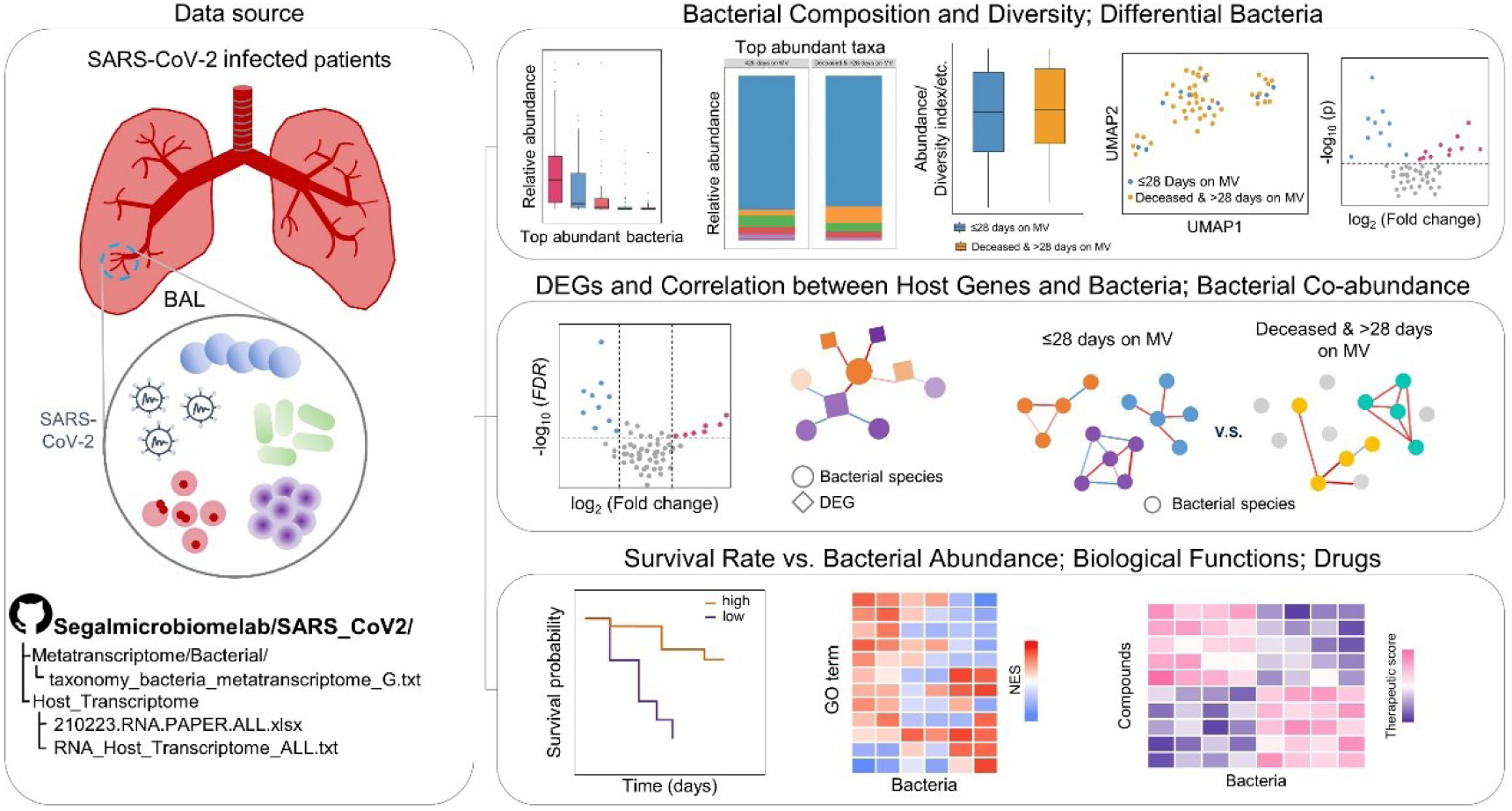
The study’s workflow The lower respiratory tract microbiome data, obtained from the BAL derived metatranscriptome and host transcriptome of severe COIVD-19 patients, were retrieved from the SARS_CoV2 repository of segalmicrobiomelab on GitHub to investigate the associations between bacterial species and host. The microbiome composition, diversity, bacterial relative abundance, bacterial co-abundance and host gene expression were compared between two clinical outcome groups (≤28 days on MV vs. Deceased & >28 days on MV). The differentially abundant bacterial species and host DEGs were further identified. SCCs were computed to determine the relationship between bacterial species and host transcriptome across all 86 BAL samples. The DEGs and SCCs were subsequently employed to identify the DEGs-associated bacterial species. Each of the differentially abundant or DEGs-associated bacterial species was later classified into high and low groups based on its bacterial relative abundance to ascertain whether the bacterial species individually were indeed linked to patient survival. The bacterial co-abundance networks and the important clusters were identified in two clinical groups separately. The functions of these bacteria were deduced and depicted in the form of heatmaps. Ultimately, the DEGs along with their positively correlated bacteria were isolated and employed in the identification of appropriate drug targets for combating COVID-19 with the impact of bacterial species.

**Table 1.** Overview of patients necessitating mechanical ventilation (MV) (N = 86)

|  | ≤28 days on<br>MV<sup>a</sup><br>(N = 21) | Deceased & >28<br>days on MV<sup>a</sup><br>(N = 65) | p-value<sup>b</sup> |
| --- | --- | --- | --- |
| Age | 65 (30 - 84) | 64 (18-87) | 0.841 |
| Gender |  |  | 0.610 |
| Female | 6 | 15 | – |
| Male | 15 | 50 | – |
| Race |  |  | 0.162 |
| African American | 0 | 8 | – |
| Asian | 0 | 3 | – |
| White | 9 | 30 | – |
| Other | 12 | 24 | – |

<sup>a</sup>MV: mechanical ventilation.
<sup>b</sup>The Mann-Whitney *U* test and the Chi-squared test were employed to compute the *p*-value for continuous and categorical data between two clinical outcome groups.

To elucidate the relationship between bacterial species and host transcriptome, SCCs were calculated across all 86 BAL samples. Following this, Fisher’s exact test was employed in conjunction with DEGs and SCCs, leading to the identification of nine DEGs-associated bacterial species, along with two additional species of questionable association. Each of the 213 differential abundant or nine plus two suspiciously DEGs-associated bacterial species was then categorized into high and low groups based on their median bacterial relative abundance to assess their individual relationships with patient survival. Subsequently, FGSEA was utilized to delineate the potential biological functions of these significant bacteria species, which were visualized as heatmaps. In the final step, DEGs along with their associated bacterial species were isolated for the identification of suitable drug targets to combat COVID-19, taking into account the influence of bacterial species in the alveolar microenvironment. Most of R scripts in this study were executed using R (version 4.2.1) on a Windows 10 desktop.

### Bacterial composition in the lower respiratory tract of critically ill COVID-19 patients

The top twenty abundant bacteria, as depicted in the two boxplots, were determined by ranking them in descending order based on their median bacterial relative abundance within each of the two clinical outcome groups separately (Figure S2). The five most abundant bacterial species identified were *Staphylococcus epidermidis*, *Mycoplasma salivarium*, *Staphylococcus aureus*, *Finegoldia magna* and *Bacillus cereus*, though their order varied slightly between groups. The stacked barplots revealed the most abundant bacterial species, genera and phyla, determined by the mean bacterial relative abundance in the deceased & >28 days on MV group at each taxonomic level in individual samples, as well as by the median bacterial abundant in two groups (Figure 2A–C and S3). The top five dominant genera were *Staphylococcus*, *Mycoplasma*, *Streptococcus*, *Prevotella* and *Enterococcus* (Figure 2B and S3B), while the most abundant phyla were *Firmicutes*, *Tenericutes*, *Proteobacteria*, *Bacteroidetes* and *Actinobacteria* (Figure 2C and S3C).

**Figure 2.**
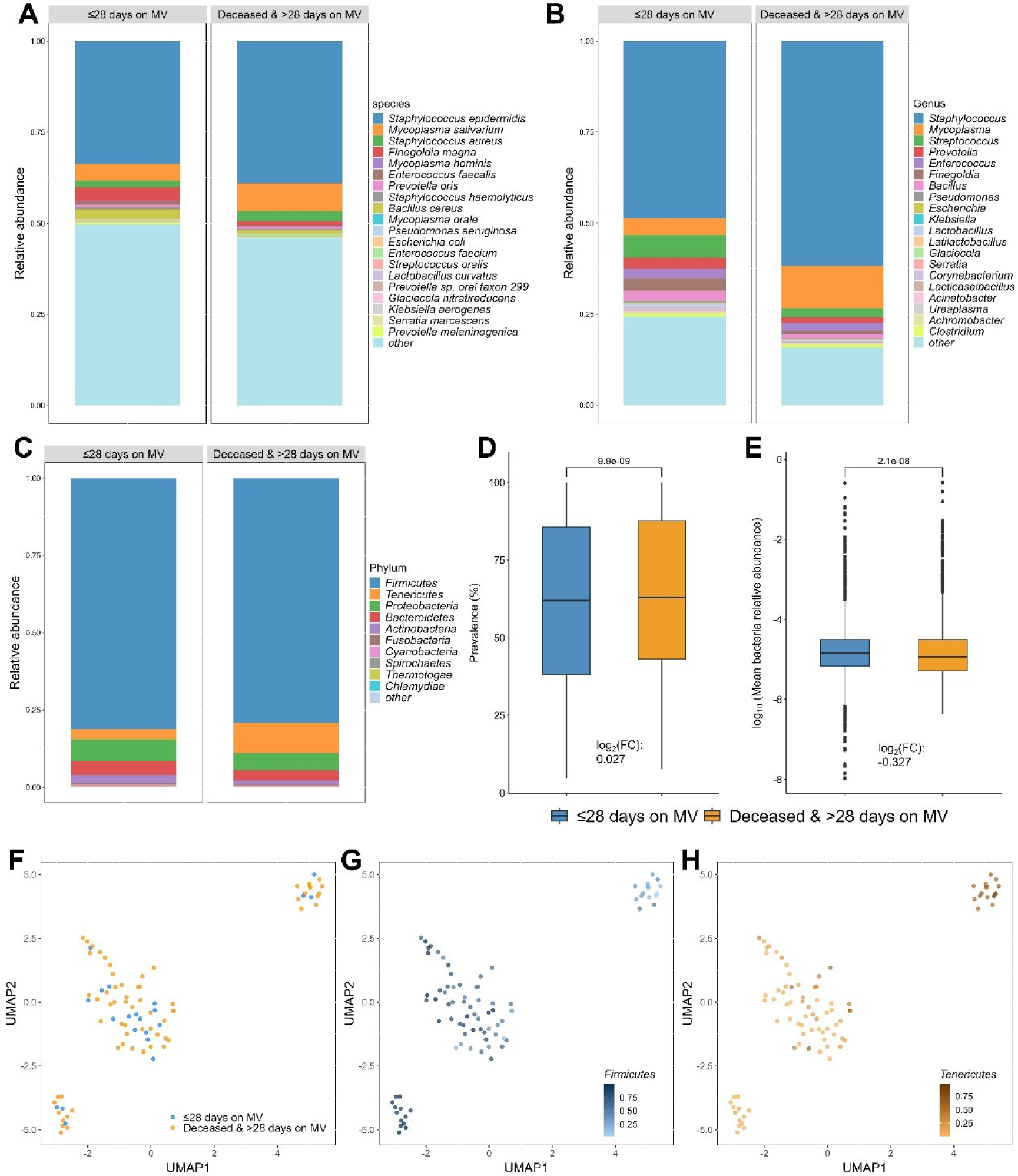
Microbial composition The top twenty or ten most abundant bacterial (A) species, (B) genera or (C) phyla, based on the mean bacterial relative abundance of corresponding taxonomic level in the deceased & >28 days on MV group, were identified and presented with their median bacterial relative abundance in the stacked barplots for two clinical groups. Figure S3 showed the bacterial relative abundance observed in each sample. (D) The prevalence of 4157 bacterial species was a little higher in deceased & >28 days on MV group. (E) On the other hand, the average relative abundance of 4157 bacteria exhibited a slight decline in the deceased & >28 days on MV group. The UMAP plots were generated using the relative abundance of bacterial species, and were color-coded to represent the (F) two clinical groups, and two of the most abundant phyla, e.g. (G) *Firmicutes* and (H) *Tenericutes*. The *p*-value was calculated using the Mann-Whitney *U* test.

The diversity indices, including the Shannon index, Richness, Simpson and Simpson reciprocal indices, as well as Pielou’s evenness, exhibited a slight decrease in the group of deceased & > 28 days on MV, although this decrease did not reach a statistical significance (Figure S4). The prevalence of 4157 bacterial species (log_2_ fold change: 0.027) and mean bacterial relative abundance (log_2_ fold change: −0.327) were showed in Figure 2D and 2E. The UMAP plot indicated that distinguishing between the two clinical groups based on the relative abundance of bacterial species was challenging, as all samples were from critically ill COVID-19 patients (Figure 2F). Further analysis identified three subgroups within the 86 BAL samples. These subgroups exhibited differing distribution of the five most abundant phyla (Figure 2G, 2H and S5A– C), with *Firmicutes* showing comparatively lower abundance in the top-right subgroup (Figure 2G) and *Tenericutes* being higher in the same subgroup (Figure 2H). The Shannon diversity index was lower in the bottom-left subgroup (Figure S5D), and the duration of ventilation days did not differ significantly among three subgroups (Figure S5E).

### Discovery of the differential abundant bacterial species between two clinical groups

We further identified a total of 213 bacterial species that exhibited differential abundance between the two groups (deceased & > 28 days on MV vs. ≤ 28 days on MV) based on the combined results of the Wilcox rank-sum test and ZicoSeq, both achieving a significance level of *p*-value < 0.05 (Figure 3A, S6 and Table S4). Figure S7 presents the first ten bacterial species found to be significantly and differentially abundant, with greater enrichment within each clinical group. Additionally, Figure S8 illustrates the inferred biological functions of these twenty bacterial species. The heatmap reveals that GO terms of these bacteria were effectively clustered according to the two clinical groups, with exceptions for *Allokutzneria albata*, *Arcobacter skirrowii*, *Psychrobacter sp. P11G3* and *Vibrio mimicus*. Genes that had positive correlations with bacteria, predominantly more abundant in the ≤ 28 days on MV group, were linked to functions such as cilium movement and the formation of microtubule bundles. Intriguingly, genes involved in these functions negatively correlated with most of the bacteria enriched in the deceased & > 28 days on MV group. Conversely, genes positively correlated with bacteria, primarily more enriched in the deceased & > 28 days on MV group, were associated with immune cells response, interleukin 6 production, phagocytosis, and so on. Genes within these functions displayed negative correlations with some of the bacteria enriched in the ≤ 28 days on MV group. Beyond these twenty bacterial species, we also pinpointed several distinctively abundant bacteria previously reported in COVID-19 related studies and in contexts involving inflammation reduction or the development of severe pneumonia and diarrhea conditions. For example, *Neisseria gonorrhoeae* and *Vibrio sp. 2521-89* were found to be more abundant in the deceased & > 28 days on MV group (*p*-value < 0.05, Figure 3B and 3C). The ≤ 28 days on MV group exhibited significantly higher abundance of *Fusobacterium periodonticum*, *Lactobacillus gasseri*, *Rhodobacter sphaeroides* and *Sorangium cellulosum* (*p*-value < 0.05, Figure 3D–G).

**Figure 3.**
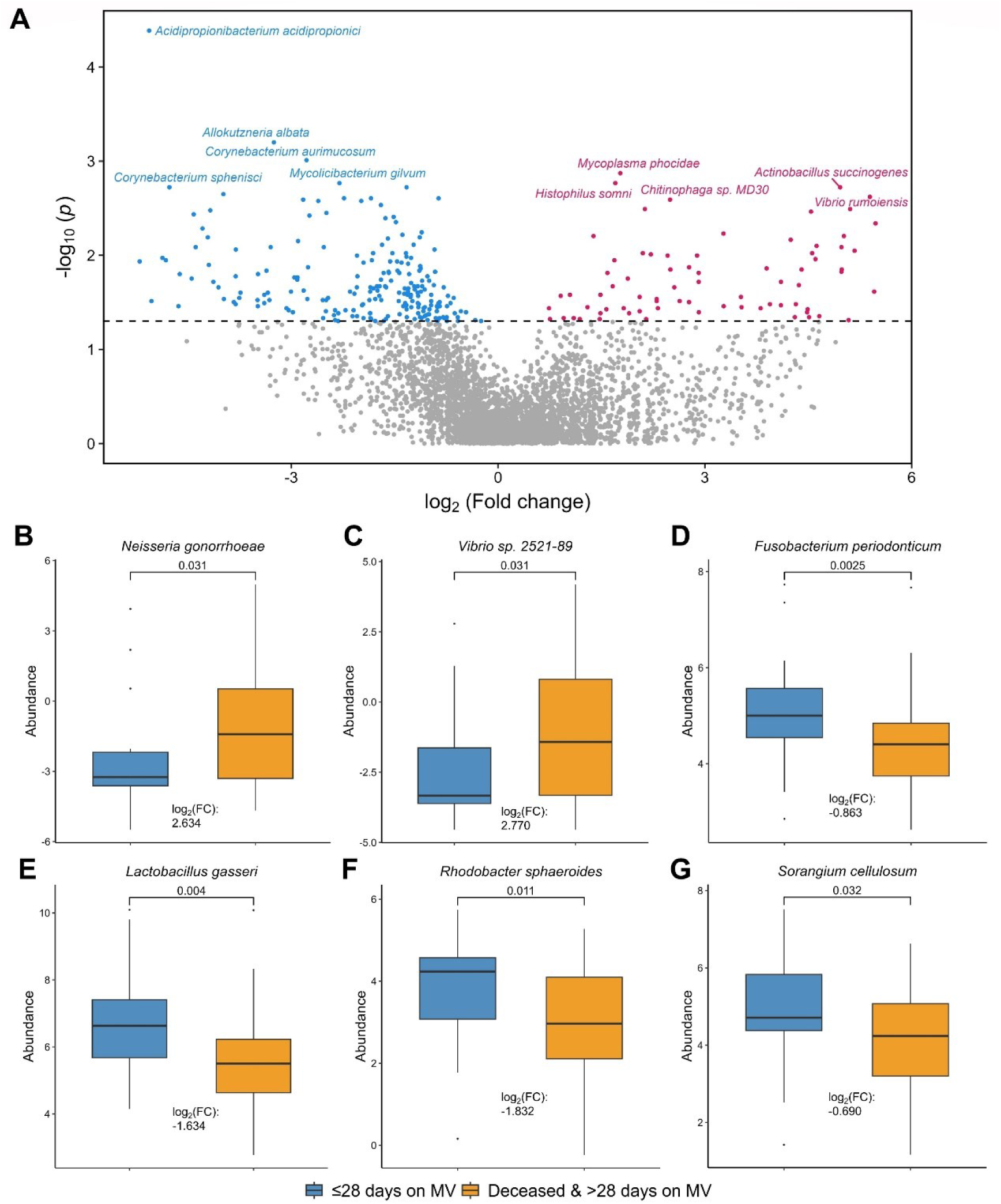
Differential abundance of bacterial species (A) A total of 284 bacterial species exhibited differential abundance, with 210 of them showing higher abundance in the ≤28 days on MV group (indicated by dark sky blue color), while 74 species were more enriched in the deceased & >28 days on MV group (indicated by dark pink color). (B) *N. gonorrhoeae* and (C) *V. sp. 2521-89* were found to be more enriched in the deceased & >28 days on MV group. On the other hand, (D) *F. periodonticum*, (E) *L. gasseri*, (F) *R. sphaeroides* and (G) *S. cellulosum* were among the bacterial species that were more abundant in the ≤28 days on MV group. The fold-change, represented as the ratio of exponential median difference between the CLR-transformed bacterial relative abundance between the two groups (deceased & >28 days on MV and ≤28 days on MV), was used to quantify the differences in abundance. The *p*-value was calculated using the Mann-Whitney *U* test. Refer to Figure S6 and Table S4 for additional information.

### Identification of the DEGs and DEGs-associated bacterial species within the host

The procedure for identifying DEGs-associated bacteria is illustrated in Figure 4A. DEGs were identified between the two groups using edgeR, resulting in the identification of a total of 141 host DEGs with a threshold of |log_2_FC| ≥1 and FDR < 0.05 (Table S5). Among these DEGs, there were 4 up-regulated genes and 137 down-regulated genes in the deceased & >28 days on MV group compared to ≤ 28 days on MV group (Figure 4B). The Spearman correlation coefficient (SCC, see Table S6) was computed to establish the associations between normalized host transcriptome and CLR-transformed bacterial relative abundance. A network was constructed using Cytoscape to illustrate the DEGs and their associated bacteria, with an absolute SCC value greater than 0.4 (Figure S9). This network included 378 edges (bacterial species and DEGs pairs), comprising 173 bacterial species and 101 DEGs. To identify bacterial species that had stronger or weaker associations with DEGs compared to non-DEGs, Fisher’s exact test was applied to each contingency table derived from the data summary for each bacterial species, with *p*-values undergoing further multiple testing correction. Ultimately, nine DEGs-associated bacterial species were identified based on the criteria of OR > 1 and FDR < 0.05. The nine DEGs-associated bacterial species were more closely correlated with DEGs than other bacteria of this network (Figure 4C and S9), as anticipated. A comprehensive list of these species is available in Table 2 & Table S7. The union of the biological functions selected from the top 20 absolute NES scores for each of the nine DEGs-associated bacterial species was visualized in a heatmap (Figure 4D). They were clustered into two subgroups and showed reverse correlation. *Blastomonas fulva* and *Sphingomonas sp. Cra20* exhibited a positive correlation with genes implicated in cilium organization, while demonstrating a negative correlation with genes linked to biological functions associated with immune responses. In contrast, the remaining seven bacterial species, namely *Borreliella valaisiana*, *Gramella forsetii*, *Hungateiclostridium thermocellum*, *Pectobacterium carotovorum*, *Providencia rustigianii*, *Raoultella ornithinolytica* and *Rickettsia canadensis*, displayed opposing results. Additionally, the bacterial species that clustered close in the heatmap also showed a greater degree of phylogenetic similarity in their 16S-rRNA genes. For example, *B. fulva* and *S. sp. Cra20*, as well as *P. carotovorum*, and *R. ornithinolytica,* exhibited close clustering in both their biological functions and the phylogenetic tree (Figure 4D and S10). Notably, *R. ornithinolytica* was the only species among the nine that also displayed a significant increase in abundance within the deceased & > 28 days on MV group (*p*-value < 0.05, Figure 4E). In addition to the nine identified DEGs-associated bacteria, we also found two additional bacteria, namely *Cedecea neteri* and *Leptospira biflexa*, which were classified as suspiciously DEGs-associated due to their Fisher FDR being less than 0.1. Furthermore, the top 25 functions for these two bacteria, arranged by absolute NES scores in descending order, were identical to those of the nine DEGs-associated bacteria (Figure 4D and S11).

**Figure 4.**
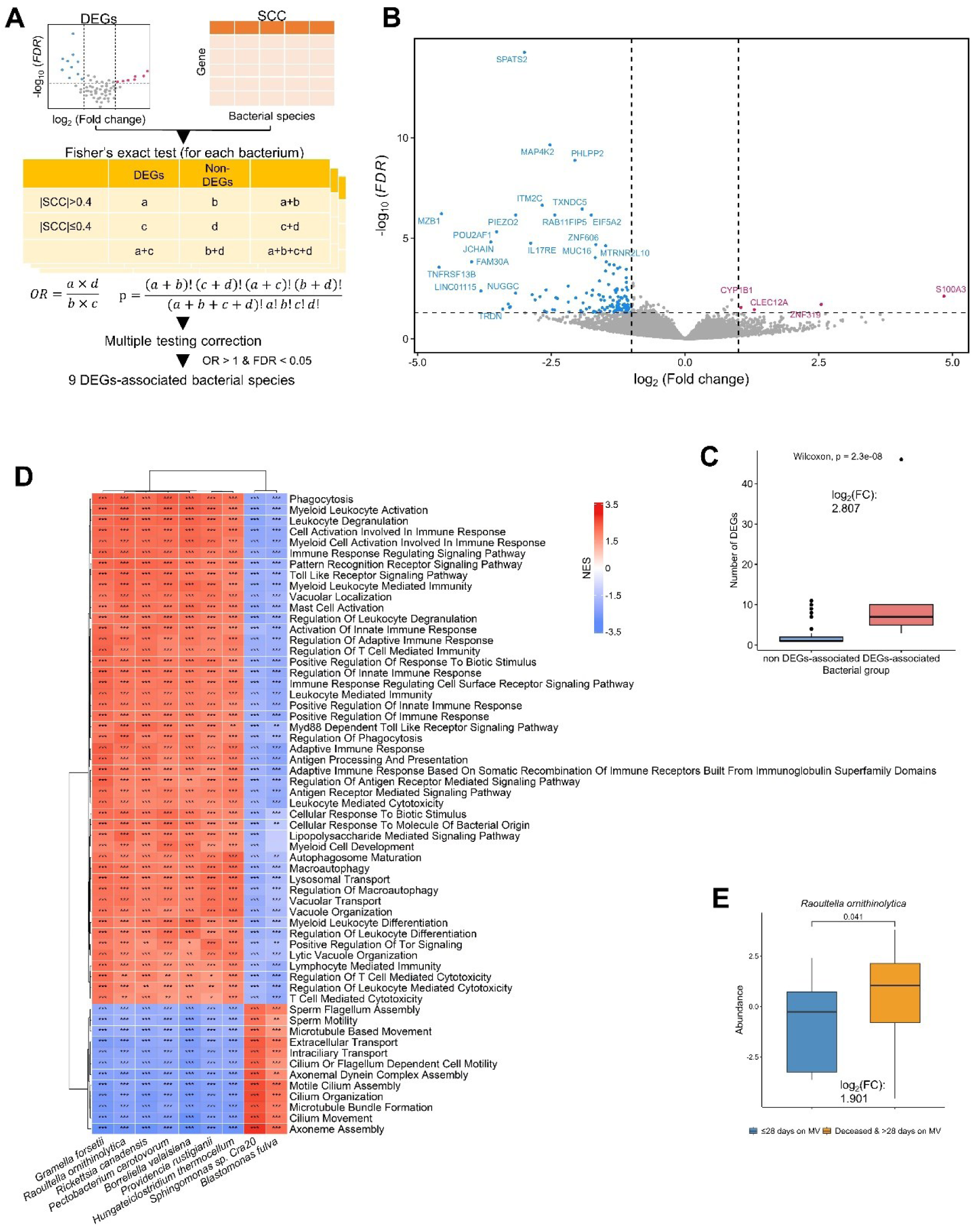
The identification of the ten DEGs-associated bacteria and their inferred biological functions (A) The flowchart showed the procedure for discerning the nine DEGs-associated bacterial species. (B) A total of 141 DEGs were discovered in the host (Table S5), with genes enriched in ≤28 days on MV group represented in dark sky blue color, while genes more abundant in the deceased & >28 days on MV group were indicated by dark pink color. (C) It was observed that the nine DEGs-associated bacterial species exhibited a higher degree of DEGs compared to the remaining bacteria within the network of DEGs and their correlated bacteria shown in Figure S9. (D) The biological functions in this heatmap were determined by combining the top twenty Gene Ontology (GO) terms with the highest absolute NES scores across the nine DEGs-associated bacteria. The GO terms of each bacterium were labeled with an asterisk to indicate their significance as follows: *: 0.01 ≤ adjusted *p*-value < 0.05; **: 0.001 ≤ adjusted *p*-value < 0.01; ***: adjusted *p*-value < 0.001. (E) The abundance of *R. ornithinolytica* significantly increased within the deceased & >28 days on MV group. The *p*-value of (C) and (E) was calculated using the Mann-Whitney *U* test.

**Table 2.**
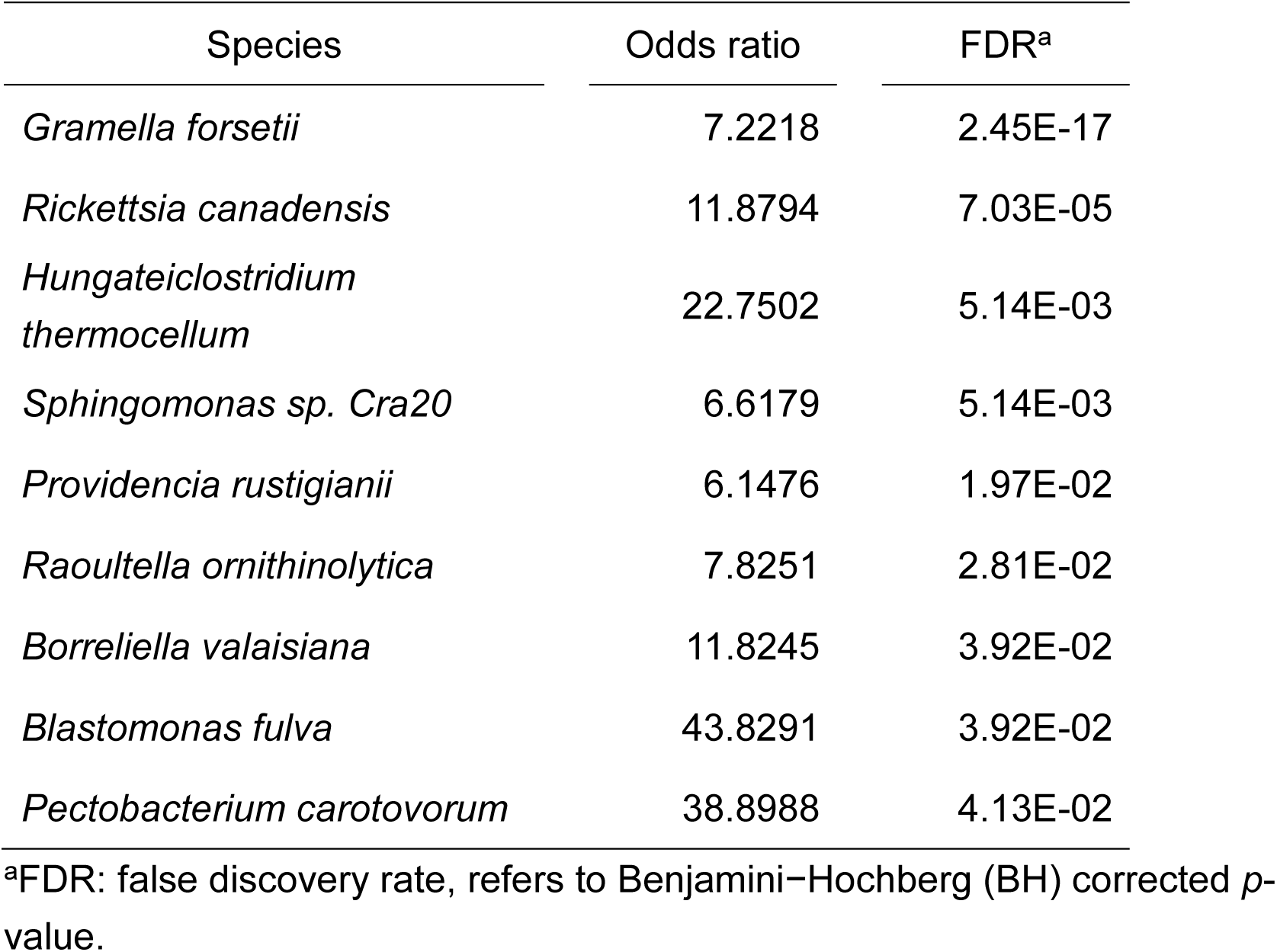
The list of DEG-associated bacteria.

### Investigation of the bacterial species related with the survival of COVID-19 patients from DEGs-associated or differential abundance of bacterial species

The 86 samples were categorized into high and low groups for each bacterial species based on the median CLR-transformed relative abundance. Specifically, patients were classified into the high group for each bacterium if their CLR-transformed relative abundance exceeded the median value; the remaining patients were classified into the low group. We then conducted a log-rank test on each of the nine DEGs-associated bacterial species, along with two potentially DEGs-associated bacterial species and those exhibiting differential abundance (*p*-value < 0.05) to assess any potential association with the survival of these severe COVID-19 patients (Table S8). A significant correlation was found between patient survival and the presence of specific bacterial species, with a total of 39 bacterial species—comprising one potentially DEGs-associated bacterium and 38 other differential bacteria— demonstrating a significant association with patient survival outcomes. Notably, patients with a lower abundance of *V. sp. 2521-89*, *Chlamydia caviae*, *Chlamydia muridarum*, *Cronobacter universalis*, *Gemmatimonas aurantiaca*, *Gordonia polyisoprenivorans*, *Salinivibrio kushneri*, *Sediminispirochaeta smaragdinae*, *Sulfuricella denitrificans*, *Waddlia chondrophila Corynebacterium atypicum*, *Microbacterium hominis* and *Plantactinospora sp. BC1* showed better survival rates (Figure 5A, S12 and S14). Boxplots derived from the differential analysis of bacterial abundance confirmed these species were present at the higher levels in the deceased & > 28 days on MV group, except for *C. atypicum*, *M. hominis* and *P. sp. BC1* (Figure 3C, S12 and S14). Conversely, patients with a higher abundance of the remaining 26 bacterial species demonstrated improved survival outcomes (Figure 5B–F, S13 and S14). The corresponding boxplots further supported these findings, indicating higher levels of these species in the ≤ 28 days on MV group, with the exception of *L. biflexa* (which is a potentially DEGs-associated but not a differential bacterium) and *Jiangella sp. DSM 45060*, which was found to be significantly more abundant in the Deceased & > 28 days on MV group (Figure 3D-G, S13 and S14).

**Figure 5.**
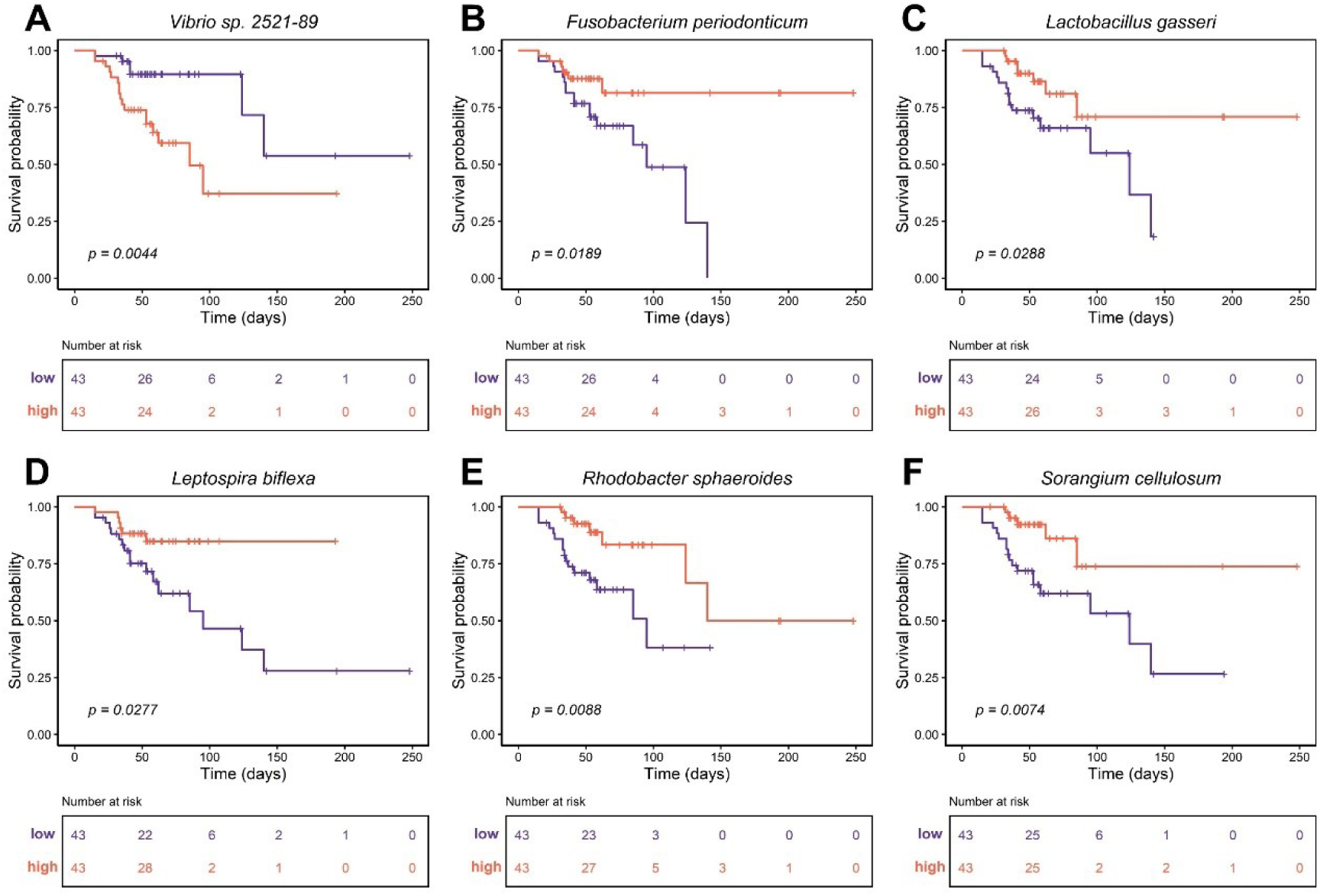
Survival related bacterial species Patients with a reduced presence of (A) *V. sp. 2521-89* demonstrated an improved survival rate. Conversely, patients who belonged to the high abundant group, including (B) *F. periodonticum*, (C) *L. biflexa*, (D) *L. gasseri*, (E) *R. sphaeroides*, and (F) *S. cellulosum*, showed a more favorable survival rate. These species were among the potentially and DEGs-associated bacterial species, and the 213 differentially abundant bacterial species. The *p*-value was computed by log-rank test (Table S8).

The biological roles of these 39 bacterial species were diverse (Figure S15) Some bacteria exhibiting higher abundance in the ≤ 28 days on MV group, and associated with better patient survival, showed positive correlations with genes involved in the cilium movement, or negative regulation of viral processes and defense responses to symbiont. Intriguingly, the genes within these GO-terms were either uncorrelated or negatively correlated with most of the bacterial species more abundant in the deceased & > 28 days on MV group, where higher abundance correlated with poorer patient survival. Conversely, some bacteria that were more abundant in the deceased & > 28 days on MV group, and associated with worse survival rate, exhibited positive correlations with genes involved in the negative regulation of interleukin 1 (beta) production, acute inflammatory response, regulation of plasma lipoprotein particle levels, etc. Notably, the genes within these GO-terms were either uncorrelated or negatively correlated with most of the bacterial species which were more abundant in the ≤ 28 days on MV group, where higher abundance was associated with better patient survival.

### Bacterial co-abundance networks in two clinical groups

To further investigate the interactions between these bacterial species, co-abundance networks were deduced for the groups of ≤ 28 days on MV and deceased & > 28 days on MV, respectively. Initially, bacterial species that were uncommon and represented as zeros in more than half of the samples were removed. Owing to the extensive number of species, only pairs of species with an absolute value of correlations ≥ 0.5 and a *p*-value ≤ 0.001 were included in each co-abundance network from different clinical outcome groups. The network for the ≤ 28 days on MV group consisted of 1158 nodes and 8253 edges, while the network for the deceased & >28 days on MV group included 1244 nodes and 5162 edges. A reduction in network complexity was observed in the bacterial co-abundance networks of two clinical groups. Topology features determined through the analyze network function in Cytoscape, including the number of edges, average number of neighbors, and network density were all lower in the deceased & >28 days on MV group (Table 3).

**Table 3.** The topology of two co-abundance networks.

| Network topology | ≤28 Days on MV<sup>a</sup> | Deceased & >28 days on MV<sup>a</sup> |
| --- | --- | --- |
| Number of nodes | 1158 | 1244 |
| Number of edges | 8253 | 5162 |
| Average number of neighbors | 14.899 | 8.848 |
| Network diameter | 13 | 14 |
| Network radius | 7 | 8 |
| Characteristic path length | 3.555 | 4.857 |
| Clustering coefficient | 0.209 | 0.232 |
| Network density | 0.014 | 0.008 |
| Network heterogeneity | 1.326 | 1.342 |
| Network centralization | 0.093 | 0.073 |
| Connected components | 26 | 41 |

<sup>a</sup>MV: mechanical ventilation.

MCODE was then used to independently identify modules with highly connected local regions, pinpointing the pivotal contributors within each complex network.^18^ Modules containing species selected by the top five abundant bacteria in the two clinical groups, the five most differentially abundant bacterial species enriched in each group, nine DEG-associated bacteria, and 39 survival-related bacteria were highlighted. In the ≤ 28 days on MV group, eight out of the 22 modules (specifically, cluster 1, 2, 3, 4, 10, 11, 13 and 17) were selected. Conversely, in the deceased & > 28 days on MV group, seven (1, 3, 4, 8, 21, 22 and 24) out of the 24 modules were isolated. To guarantee that the nodes were aligned consistently across both networks, we combined the nodes and edges from both networks (Table S9), which served as the basis for the structure and were visualized using Cytoscape. Figure 6 demonstrates that the bacterial co-abundance networks showed differences between the two clinical groups, with bacterial species sharing similar phylogenetic and taxonomy categories tending to be co-expressed together (Figure S16).

**Figure 6.**
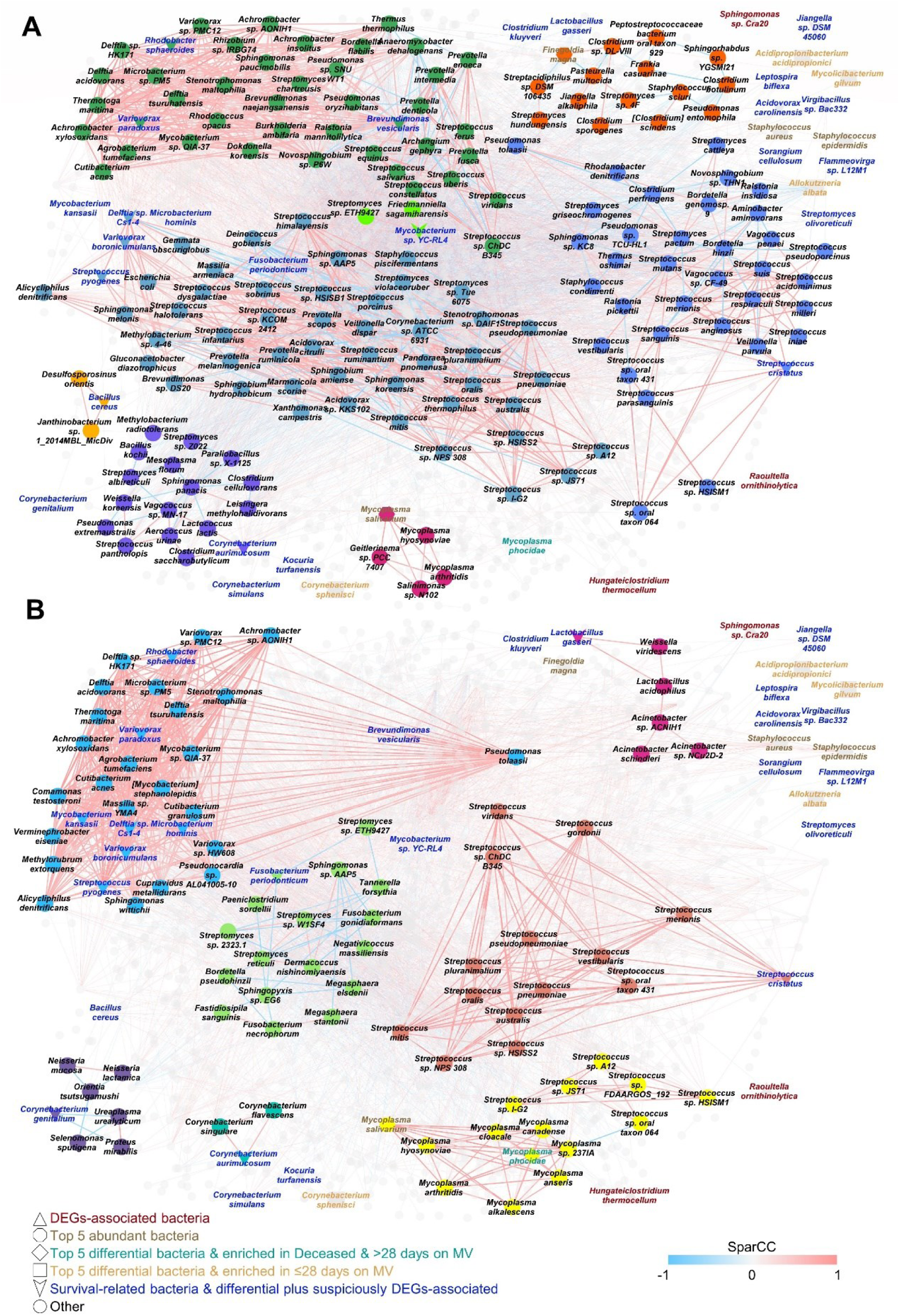
The bacterial co-abundance networks in two clinical groups. MCODE generated clusters that consisted of the five most notable and distinctively abundant bacterial species, DEGs-associated bacteria, the top five abundant bacteria in the two clinical groups, or 39 survival-related bacteria were highlighted and illustrated separately in the groups of (A) ≤28 Days on MV and (B) deceased & >28 Days on MV. See also Table S9.

### Drug discovery targeting genes correlated with bacterial species

To identify potential compounds for combating COVID-19 influenced by bacterial presence, we applied a gene-expression-based methodology developed in our research.^20^ This study evaluated the therapeutic potential of compounds to alter gene expression profiles related to each bacterium. Our approach is based on the hypothesis that enhancing gene expression associated with beneficial bacteria while inhibiting expression linked to harmful bacteria could offer a novel treatment strategy for COVID-19. Here, “beneficial” bacteria are defined as those predominantly found in the high survival probability subgroup and enriched in patients who required MV for ≤ 28 days. Conversely, “harmful” bacteria refer to those found in the low survival probability subgroup and enriched in deceased patients or those with > 28 days on MV. In the analysis conducted at 6 hours post-drug exposure model (d6), palbociclib showed a significantly high therapeutic score in beneficial bacteria and a low score in harmful bacteria, indicating its potential to positively influence gene expression profiles associated with beneficial bacteria and to negatively affect those associated with harmful bacteria (Figure 7A). Similarly, compounds like QL-X-138 (PubChem CID: 73707530), crizotinib and XMD-1499 (PubChem CID: 73265211) demonstrated perturbation capabilities akin to palbociclib. In the 24-hour post-drug exposure model (d24), several compounds, including RS-17053 (PubChem CID: 3894573), RS-504393 (PubChem CID: 9953769), barasertib, fulvestrant, duloxetine, NNC-05-2090 (PubChem CID: 9888030), terfenadine and amoxapine, exhibited significantly high therapeutic scores for beneficial bacteria and low scores for harmful bacteria (Figure 7B). The complete drug discovery results have been visualized using a heatmap with hierarchical clustering to highlight data patterns (Figure S17). These results highlight the potential of compounds that can differentially influence the gene expression profiles associated with beneficial and harmful bacteria as viable options for COVID-19 treatment strategies.

**Figure 7.**
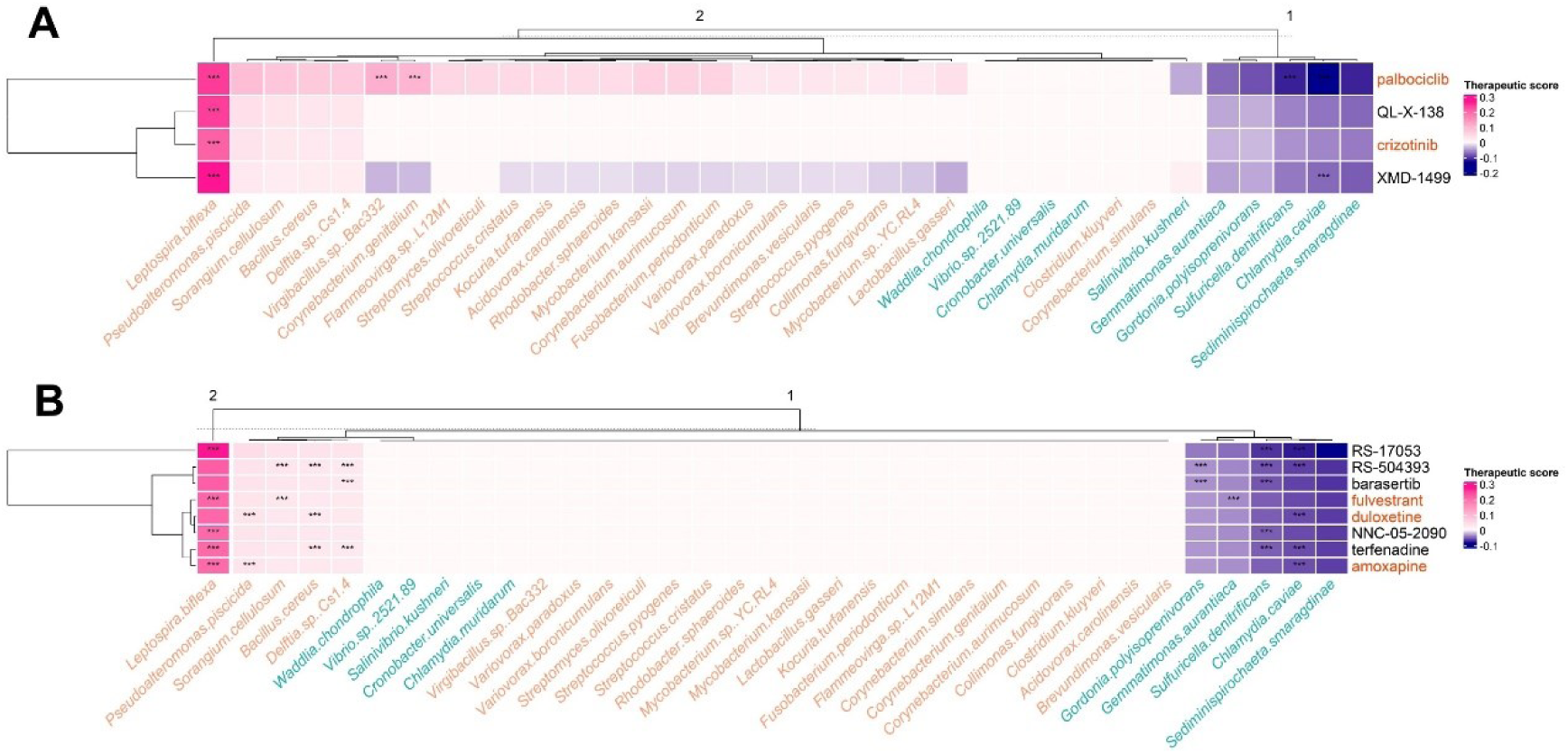
Therapeutic score of highlighted compounds in our gene-expression-based drug prediction Hierarchical clustering reflects the similarity of ability for gene perturbation between compounds and the relevance of bacterium-related gene expression between bacteria in (A) drugs for 6 hours (d6) prediction model (B) drugs for 24 hours (d24) prediction model. The rows represent compounds in the prediction. FDA-approved compounds are labeled in red. Compounds with zero value of sum therapeutic score across bacteria have been removed from this heatmap. The columns represent different bacteria-related expression profiles. Beneficial bacterium is labeled in orange; harmful bacterium is labeled in blue. The highest therapeutic score is labeled in deep pink; the lowest therapeutic score is labeled in dark blue. The significant of the prediction is marked with *: 0.01 ≤ *p*-value < 0.05; **: 0.001 ≤ *p*-value < 0.01; ***: *p*-value < 0.001. Refer to Figure S17 for the therapeutic score of complete compounds.

## DISCUSSION

There have been limited investigations into the direct sampling of the microbiome in the lung or lower respiratory tract of COVID-19 patients since the onset of the pandemic. Instead, most studies have focused on collecting samples from the upper respiratory tract through nasopharyngeal swabs. However, the primary location of disease activity for SARS-CoV-2 is the lower respiratory tract, specifically within the alveolar space. Bronchoscopy, coupled with Bronchoalveolar Lavage (BAL), is a minimally invasive procedure utilized to collect samples from a lung subsegment.^21^ During this procedure, sterile normal saline is introduced into the subregion of the lung, then suctioned out, and the fluid is collected for analysis.^21^ Sulaiman and his team have utilized this technique to gather data on the microbiome characteristics in the respiratory system of severely ill COVID-19 patients.^16^ To explore the role of bacterial species in influencing the duration and of MV usage and the severity of COVID-19 patients, we accessed data from Sulaiman’s study available on the GitHub repository. This data was then filtered and categorized the samples into two distinct clinical group: the “≤ 28 Days on MV” and the “deceased & > 28 Days on MV” groups.

In both clinical groups, the five most abundant bacterial species identified were *S. epidermidis*, *M. salivarium*, *S. aureus*, *F. magna* and *B. cereus* (Figure 2A, S2 and 3A). These bacteria have been implicated in airway infections and inflammation in previous reports. Kim et al. provided evidence that *S. epidermidis*, a common colonizer in the human mucosa including the respiratory tract, mediates frontline antiviral protection against influenza A virus (IAV) by interferon (IFN)-λ-dependent innate immune mechanisms.^22^ *M. salivarium* has been associated with poor clinical outcome in the lower respiratory tract of mechanically ventilated COVID-19 patients and in cases of ventilator-acquired pneumonia.^10,16^ *S. aureus* pneumonia is known as a common cause of hospital-acquired pneumonia in patients on ventilators.^11^ *F. magna* has been identified as the most common anaerobic isolate in patients with chronic nasal inflammatory diseases.^23^ Although *B. cereus,* primarily known for causing food poisoning, is a rare cause of lower respiratory tract infections, instances of *B. cereus* pneumonia can be fatal, indicating its potential as a pathogenic threat.^24,25^ Few differences were observed in the prevalence and mean bacterial relative abundance between two groups (Figure 2D and 2E). The diversity indices experienced a slight decrease in the deceased & > 28 Days on MV group, yet this change did not reach statistical significance (Figure S5). This pattern may reflect the common critical condition of COVID-19 patients in both groups.

A total of 213 bacterial species exhibited differential abundance between the two groups (Figure 3, S6 and Table S4). Among these, specific bacteria previously identified in COVID-19 research and in contexts associated with inflammation reduction, severe pneumonia or diarrhea were recognized. *N. gonorrhoeae*, though rarely associated with unilateral lobar pneumonia^26^, was found to be significantly more abundant in the deceased & >28 days on MV group (Figure 3B), aligning with some COVID-19 cases presenting unilateral lobar pneumonia.^27^ Pathogenic *Vibrio* species can lead to gastroenteritis, which is marked by symptoms such as diarrhea, vomiting, fever and so on, often resulting from the consumption of contaminated fish and seafood.^28–31^ In addition to *V. sp. 2521-89,* which shares a close genetic relationship with *V. cholerae*, our findings indicated that *V. mimicus*, *V. rotiferianus*, and *V. rumoiensis* were also significantly more enriched in the Deceased & >28 days on MV group (Figure 3C and Table S4). Nardelli et al.^32^ observed a significant reduction in *F. periodonticum* among the species detected in bacterial 16S rRNA sequencing of nasopharyngeal swabs from COVID-19 patients compared to control, with an even greater decrease in the deceased & > 28 days on MV group (Figure 3D). Research by Vaezi et al.^33^ found a significant reduction in IL-6 serum levels in COVID-19 patients administered a synbiotic capsule containing prebiotics and multi-strain probiotics, including *L. gasseri*, compared to the placebo group, with *L. gasseri* also reduced in abundance in the deceased & >28 days on MV group (Figure 3E). Abdelmageed et al.^34^ demonstrated the anti-inflammatory potential of lipopolysaccharide from *R. sphaeroides* (LPS-RS) in rat models of acute lung injury, with our findings showing a significant reduction in *R. sphaeroide*s in the deceased & >28 days on MV group (Figure 3F). *S. cellulosum* is capable of producing Carolacton, a *MTHFD1* inhibitor, which demonstrates inhibitory effects against a range of RNA viruses, including SARS-CoV-2.^35,36^ Intriguingly, we found that *S. cellulosum* reduced in the deceased & > 28 days on MV group (Figure 3G).

The inferred biological functions associated with the ten differential bacteria, which were enriched in either of the two groups, revealed that the top ten bacteria that were significantly more abundant in the deceased & > 28 days on MV group exhibited a greater number of GO terms positively enriched in immune responses (Figure S7 and S8). The identified functions encompassed the acute inflammatory response, a range of cytokine-related activities, including the production of IL-1, IL-6, IL-8, and interferon (IFN)-γ. Additionally, functions encompassed phagocytosis, chemotaxis and the migration of phagocytes, as well as the positive regulation of CD4^+^ alpha-beta T cell differentiation, among other processes. While these results indicate robust immune activation in the deceased & > 28 days on MV group, numerous studies on COVID-19 have demonstrated that stronger immune responses do not necessarily correlate with improved survival outcomes.^37–42^ In fact, the relationship between immune response and patient survival appears to be nonlinear, with both insufficient and excessive immune activation potentially leading to poor outcomes. For instance, research has shown that precise calibration of IFN-I response is crucial for patient survival, as both insufficient and excessive IFN activation can be life-threatening.^37,38^ This concept is further supported by Del Valle et al., who revealed an inverse correlation between inflammatory markers (IL-6, IL-8, tumor necrosis factor TNF-α) and survival rates in COVID-19 patients.^39^ In non-survivors, levels of acute-phase proteins such as C-reactive protein (CRP), ferritin, serum amyloid A (SAA), procalcitonin, and IL-6 exhibited an upward trend, while in survivors, these markers either remained stable or showed a decline.^40,41^ These findings align with our observations, where the deceased & > 28 days on MV group showed enhanced immune-related functions, suggesting that excessive immune activation may contribute to poorer outcomes rather than promoting survival.

In addition to performing a differential analysis of bacterial species, we applied Fisher’s exact test to explore the host DEGs and SCCs between bacteria and host gene. This analysis led to the identification of nine DEGs-associated bacterial species (Table 2). Figure S9 illustrates the network between host DEGs and their correlated bacterial species, with absolute SCC values exceeding 0.4. The analysis revealed that these nine DEGs-associated bacterial species had a higher degree of correlated DEGs compared to other bacteria in this network (Figure 4C). Additionally, most of these DEGs-associated bacteria were found to positively associate with the host immune response, yet negatively correlate with cilium functions (Figure 4D). Moreover, the bacterial species that clustered more closely in the heatmap showed greater genetic similarity, particularly in their 16S-rRNA genes. For example, *B. fulva* and *S. sp. Cra20*, as well as *P. carotovorum*, and *R. ornithinolytica*, exhibited close clustering in both their inferred functions and the evolutionary tree (Figure 4D and S10). Among these nine species, only *R. ornithinolytica* significantly increased within the deceased & >28 days on MV group (*p*-value < 0.05, Figure 4E). *R. ornithinolytica* is a newly recognized bacterium associated with human infections.^43^

We further identified 39 bacterial species associated with patient survival from the potentially and DEGs-associated or differentially abundant bacterial species. Specifically, *V. sp. 2521-89*, *C. caviae*, *C. muridarum*, *C. universalis*, *G. aurantiaca*, *G. polyisoprenivorans*, *S. kushneri*, *S. smaragdinae*, *S. denitrificans* and *W. chondrophila* were found in higher abundances in the deceased & >28 days on MV group (Figure 3C and S12), with lower abundant groups of these microorganisms experiencing more favorable survival outcomes in severe COVID-19 cases (Figure 5A and S12), classifying them as detrimental species. Conversely, patients with a higher abundance of the 25 bacterial species showed better survival rates compared to those with low abundances of these bacteria (Figure 5B–F and S13). These bacterial species, except *L. biflexa* (which is a potentially DEGs-associated but not a differential bacterium), were also more abundant in the ≤ 28 Days on MV group (Figure 3D–G and S13), suggesting that these bacteria positively influence the survival of severe COVID-19 patients. Thus, they are considered as potentially beneficial organisms in the context of severe COVID-19. Four species, *C. atypicum*, *J. sp. DSM 45060*, *M. hominis* and *P. sp. BC1* exhibited ambiguous characteristics (Figure S14). *C. atypicum, M. hominis and P. sp. BC1* were more abundant in the ≤ 28 Days on MV group, yet higher abundance of *C. atypicum, M. hominis and P. sp. BC1* correlated with poorer survival rates. *J. sp. DSM 45060*.was more enriched in the deceased & > 28 days on MV group, but a higher abundance of this bacterium was associated with better survival rates, indicating complex relationships between bacterial presence and patient outcomes.

A loss of network complexity was evident in the bacterial co-abundance networks of two clinical groups. Topology characteristics, such as the number of edges, average number of neighbors, and network density were all reduced in the deceased & >28 days on MV group (Table 3). This finding aligns with Hernández-Terán et al.^44^ who observed a decrease in structural complexity from mild to fatal COVID-19 cases. To enhance the identification of significant bacteria and streamline the complex network representation, we utilized MCODE-generated clusters comprising the top five abundant bacteria in the two clinical groups, the five most differentially abundant bacterial species enriched in each group, nine DEG-associated bacteria, and 39 survival-related bacteria. These clusters were presented in Figure 6, showing differences in the bacterial co-abundance networks between the two clinical groups. For example, species within the *Streptococcus* genera appeared to coexist independently in each COVID-19 group. Moreover, certain bacterial species that fall within comparable phylogenetic and taxonomic classifications exhibited co-expression patterns (Figure 6, S16 and Table S9). Notably, a survival-related bacterium, *Delftia sp. Cs1-4*, and its co-expressed species *Acidovorax citrulli, Alicycliphilus denitrificans*, *Comamonas testosteroni*, *Variovorax boronicumulans*, and *Verminephrobacter eiseniae*, all belonging to the family *Comamonadaceae*, demonstrated co-expression in clusters of the ≤ 28 Days on MV group (designated as cluster 2, located at the center of Figure 6A) and the deceased & > 28 Days on MV group (identified as cluster 1, situated in the top-left section of Figure 6B).

In our drug discovery process, we utilized a gene-expression-based approach capable of evaluating the gene perturbation effects of compounds across various cell types.^20^ This approach enables the identification of compounds that excel in enhancing the expression profile of beneficial bacteria while suppressing those of harmful bacteria. From d6 model results, palbociclib and XMD-1499 were found to significantly enhance the expression profile of beneficial bacteria and suppress that of harmful bacteria, highlighting their potential in COVID-19 treatment (Figure 7A). Palbociclib, an FDA-approved CDK4/6 inhibitor for breast cancer treatment,^45^ has been suggested to have a short-term immunosuppressive effect that might delay COVID-19 progression.^46^ Yet, another study proposes that palbociclib could facilitate SARS-CoV-2 cell entry by reducing ACE2 degradation,^47^ presenting a complex picture that warrants further research into its effects on COVID-19. Despite the promising gene perturbation ability shown by XMD-1499 in our analysis, there’s a lack of research regarding its use in COVID-19 treatment. QL-X-138, known as a Bruton’s tyrosine kinase/Mitogen-activated protein kinase interacting kinase inhibitor for lymphoma and leukemia,^48^ showed a high therapeutic score for promoting the expression of beneficial bacteria. This finding suggests its potential utility against COVID-19, with speculation that QL-X-138 may inhibit SARS-CoV-2 infection and inflammation, though this also requires further investigation.^49^ For Crizotinib, crizotinib exhibits anti-bacterial activity against Gram-positive bacteria. In addition, crizotinib treatment reduces pulmonary inflammation and increase survival rate in mice model.^50^

In the d24 model findings, several compounds were pinpointed as potential agents to promote the expression of genes associated with beneficial bacteria while inhibiting those linked to harmful bacteria, specifically RS-17053, RS-504393, barasertib, fulvestrant, duloxetine, NNC-05-2090, terfenadine and amoxapine (Figure 7B). RS-17053 exhibits a preference for the α1a-adrenoceptor subtype over the α1b- and α1d-adrenoceptor subtypes,^51^ yet its role in COVID-19 management remains to be explored. RS-504393, a CCR antagonist, has been studied in lung injury models^52^ and has demonstrated binding affinity to the SARS-CoV-2 spike protein through molecular docking simulations, suggesting a potential antiviral effect against COVID-19.^53^ Limited research exists on the effects of barasertib, fulvestrant, and NNC-05-2090 on COVID-19. Although duloxetine primarily acts as an FDA-approved antidepressant, studies have shown that it can alter bacterial community composition.^54^ Moreover, in mice model, alpha diversity and beta diversity in gut microbiome are altered after duloxetine treatment.^55^ Terfenadine, an antihistamine, is considered to potentially inhibit SARS-CoV-2 endocytosis, though this effect requires further validation.^56^ In the concept of bacterial composition alteration, terfenadine increase *Salmonella Typhimurium* loads in a synthetic gut community with twenty commensals under high drug concentration.^57^ In the general condition, amoxapine acts as an antidepressant.^58^ Interestingly, in research on bacterial control, the antimicrobial effects of amoxapine have been investigated, highlighting its potential to influence bacterial community composition.^59^ In another research, amoxapine attenuates the effect of mycophenolate mofetil, which might cause enteropathy, on some bacteria abundance, such as *Gammaproteobacteria*.^60^

Overall, most of the highlighted compounds exhibit the ability to alter bacterial composition or demonstrate antiviral effects against SARS-CoV-2. These properties may explain their potential benefits in improving COVID-19 outcomes, possibly through the modulation of bacterial composition. Consequently, compounds such as palbociclib, XMD-1499, QL-X-138, RS-17053, RS-504393 and duloxetine are advocated for further study as prospective COVID-19 therapeutics.

In summary, the microbiome composition among critically ill COVID-19 patients varies, particularly between the two groups categorized as the ≤ 28 days on MV and the deceased & >28 days on MV. Our study revealed that 213 bacteria species showed significant and differential abundance levels between these groups. Additionally, we identified nine plus two DEGs-associated bacterial species. Notably, within these differentially abundant or potentially and DEGs-associated bacterial species, we found 39 that are closely linked to patient survival. We also elucidated the biological functions of these bacteria interest. Our analyses further indicated a reduction in network complexity in the deceased & > 28 days on MV group. Importantly, DEGs showing a positive correlation with the 35 survival-related bacteria emerge as potential targets for drug repurposing in COVID-19 treatment strategies.

### Limitations of the study

We applied the CLR-transformation to adjust the bacterial relative abundance, aiming to reduce data skewness and compositionality bias. While we did not assess other transformed methodologies, we incorporated the ZicoSeq approach in identifying the differential bacterial species. The insufficiently strong correlation between bacterial abundance and the host DEGs may diminish the efficacy of drug prediction methods in determining the microbiome’s influence on therapeutic interventions. Furthermore, the results of these predicted drugs require additional comprehensive experimental validation.

## RESOURCE AVAILABILITY

### Lead contact

Further information and requests for resources and reagents should be directed to and will be fulfilled by the lead contact, Hsueh-Fen Juan.

### Materials availability

This study did not generate new unique reagents.

### Data and code availability

- This paper does not report original code.
- Any additional information required to reanalyze the data reported in this paper is available from the lead contact upon request.

## ACKNOWLEDGMENTS

We thank Professor Hirotada Mori from Guangdong Academy of Agricultural Sciences for the useful discussion. This work was supported by National Science and Technology Council (NSTC 109-2221-E-002-161-MY3, NSTC 109-2221-E-010-011-MY3, NSTC 112-2221-E-A49-061-MY3, NSTC 112-2320-B-002-022, and NSTC 113-2221-E-002-149-MY3), Research Proposal for NTU Core Consortiums (NTU-113L8503) and the Center for Advanced Computing and Imaging in Biomedicine (NTU-113L900701 and NTU114L900701) from The Featured Areas Research Center Program within the framework of the Higher Education Sprout Project by the Ministry of Education (MOE) in Taiwan. The funding agencies had no role in the design, analysis, and interpretation of the data or writing of the manuscript.

## AUTHOR CONTRIBUTIONS

H.-C.H. and H.-F.J. conceived and supervised the work. K.-P.C., C.-H.H. and K.-Y.C. analyzed the data. C.-L.H. helped with data analysis. K.-P.C., C.-H.H., K.-Y.C., H.-C.H. and H.-F.J. interpreted the results and wrote the manuscript. All authors read and approved the final manuscript.

## DECLARATION OF INTERESTS

The authors declare no competing interests.

## SUPPLEMENTAL INFORMATION

**Document S1. Figures S1–S17** (this is the main PDF)

**Document S2. Tables S1–S10** (this is the link for the Tables S1–S10 that have been privately deposited on the Figshare repository)

## STAR★METHODS

### KEY RESOURCES TABLE

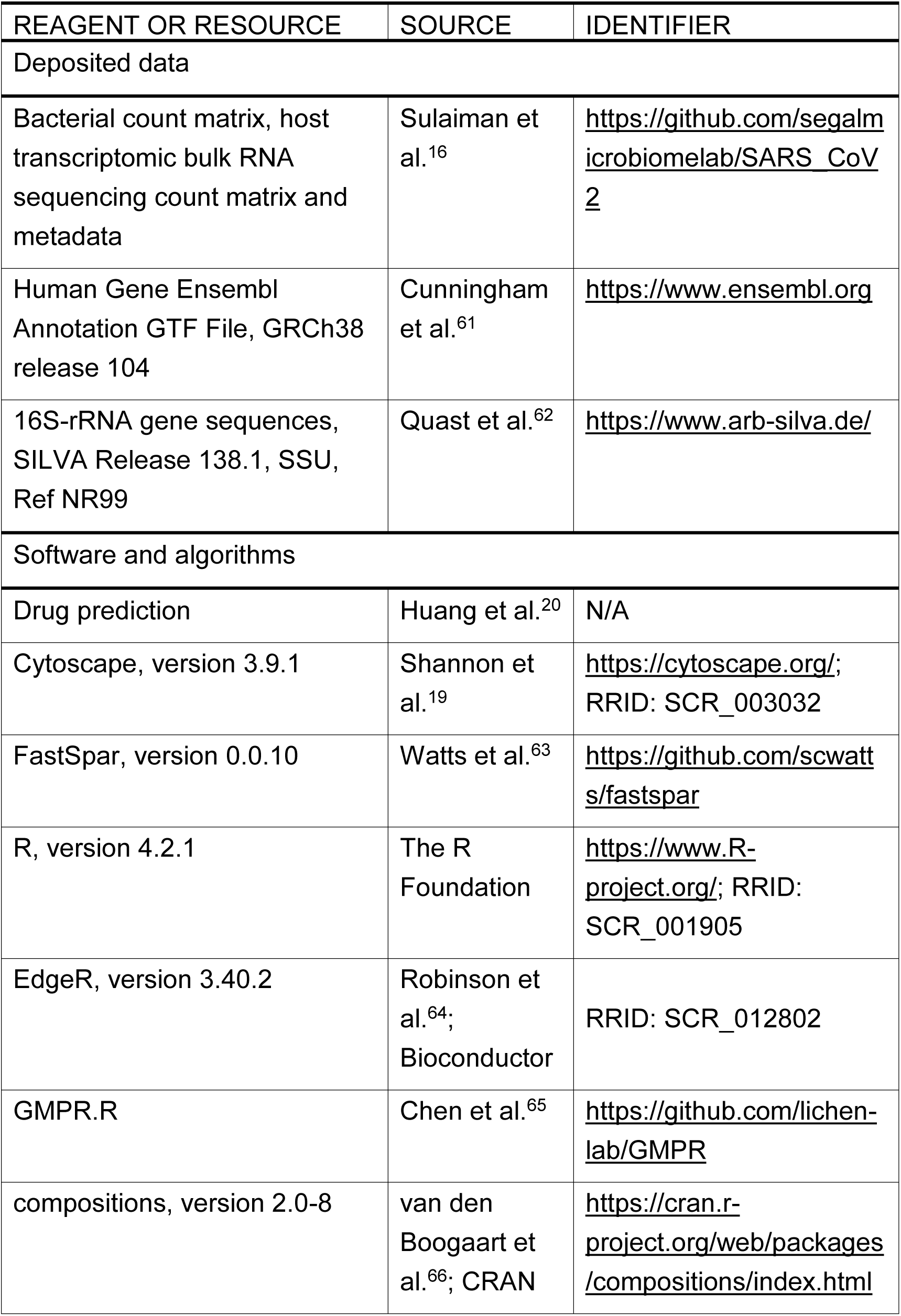
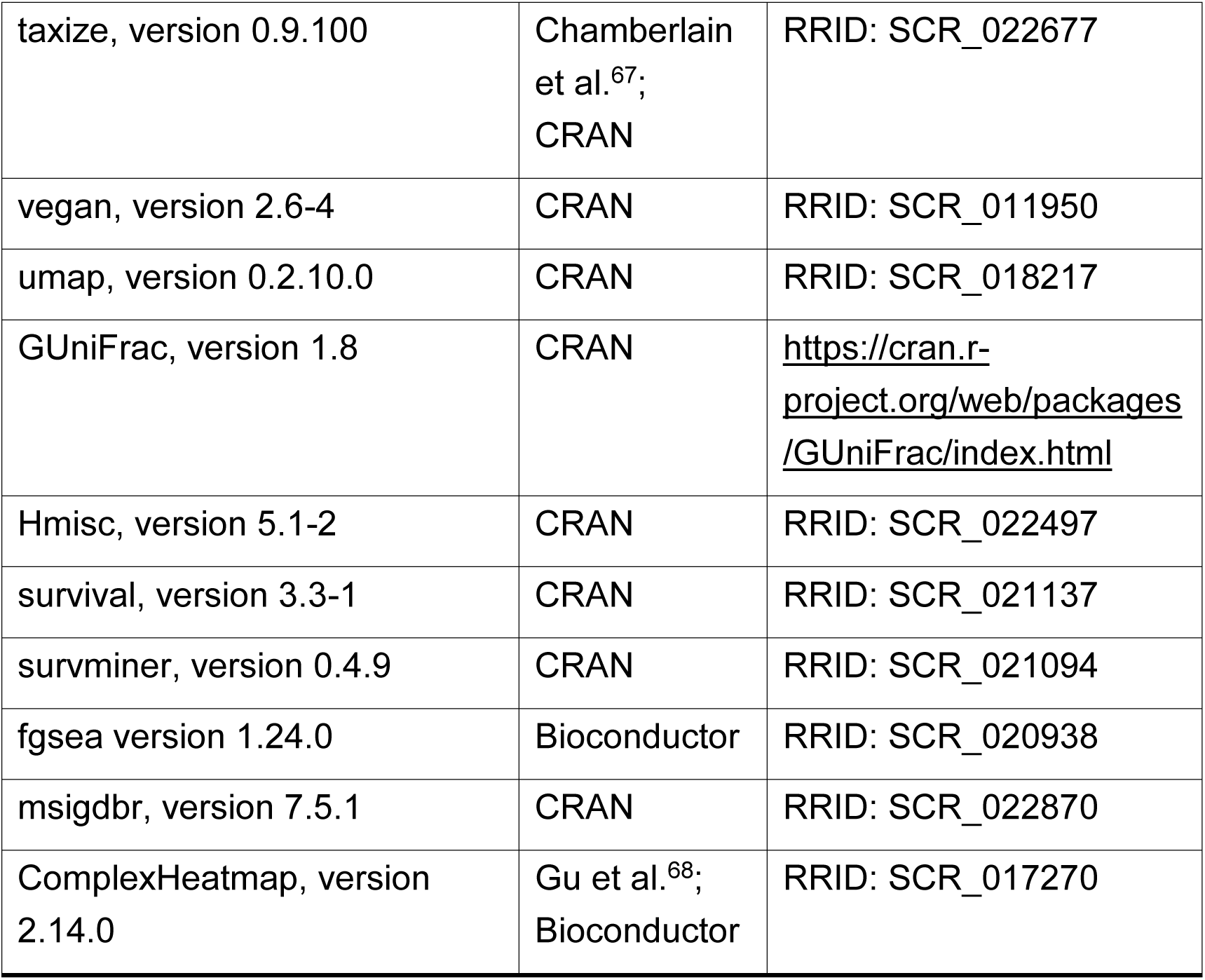

### EXPERIMENTAL MODEL AND STUDY PARTICIPANT DETAILS

The study cohort comprised 86 severely ill COVID-19 patients, divided into two clinical groups based on the host transcriptome: “≤ 28 days on MV” (*N* = 21) and “Deceased & >28 days on MV” (*N* = 65). We utilized data files containing bacterial count, metadata and host transcriptome matrices from the segalmicrobiomelab’s GitHub repository.^16^

## METHOD DETAILS

### Data sources and data pre-processing

The metadata, bacterial count matrix, and host transcriptomic bulk RNA sequencing count matrix (210223.RNA.PAPER.ALL.xlsx, taxonomy_bacteria_metatranscriptome_G.txt and RNA_Host_Transcriptome_ALL.txt) were obtained from the segalmicrobiomelabs’s SARS_CoV2 GitHub repository.^16^ The Human Gene Ensembl Annotation GTF File (Homo_sapiens.GRCh38.104.gtf.gz, version: GRCh38 Release 104) was downloaded on November 9, 2021.^61^ The GTF file was used to filter for genes that are either lncRNAs or protein-coding genes, while other biotypes were excluded from this study.

Because the primary site of SARS-CoV-2 disease activity is the lower respiratory tract, only lower airway (BAL, *N* = 122) specimens were used. The host transcriptome was filtered to exclude gene symbols not classified as lncRNAs or protein-coding genes, as well as expressions accounting for less than 20% of the BAL samples. A boxplot was utilized to visualize gene expression in the BAL samples. The 36 samples with skewed gene distribution/low abundance or lacking metadata information were discarded. We then continued to use EdgeR (version 3.40.2)^64^ to acquire the normalized counts per million (CPM) with a log_2_ transformation (Table S2).

Bacterial species with prevalence (raw count > 0) in less than 20% of the BAL samples were removed. Subsequently, only those 86 samples (Table S1) for which the host transcriptome matrix remained were retained. GMPR was then employed in this filtered bacterial raw count matrix for normalization across samples, using the default settings.^65^ The normalized count matrix was subsequently transformed into a matrix reflecting the relative abundance of bacteria, e.g. the normalized counts for each bacterium were divided by the total normalized counts within individual samples. The center log ratio (CLR) transformation was applied to mitigate data skewness and compositional bias in bacterial relative abundances.^69^ To prevent infinite or NaN values during the CLR transformation, a nonzero minimal value of 5.602744e-08 was added. The CLR transformation was performed using the R package compositions (version 2.0-8, Table S3).^66,70^ The taxonomic levels of these bacterial species were retrieved from the NCBI database on February 9, 2022, using the R package taxize (version 0.9.100; R v4.0.3, Ubuntu 18.04.6 LTS machine).^67^ The bacterial relative abundances of genus and phylum levels were summed for each taxon from the processed bacterial relative abundance matrix and the taxa classified as NA were removed.

### Composition, diversity and differential abundance of bacteria in the lower respiratory tract

The twenty most abundant bacterial species and genera, along with ten most abundant phyla, based on the median or mean bacterial relative abundance at each taxonomic level, were identified. These were presented in boxplot and stacked barplot, either across all 86 BAL samples or within two classified clinical groups. The remaining bacterial taxa, beyond the top abundant ones, were aggregated and categorized into the ‘other’ category in each sample or group in the stacked barplot.

The indices of Shannon (*H*^′^), Simpson (*D*), Inverse Simpson (1/D) and Pielou’s evenness (*J*^′^)^71–75^ were calculated using the R package vegan (version 2.6-4)^76^ as follow:

- Shannon index: *H′ = −∑^s^_i=1_p_i_ ln p_i_*
- Simpson index: *D = 1 − ∑^s^_i=1_ p^2^_i_*
- Inverse Simpson index: *1/D = 1/∑^s^_i=1_ p^2^_i_*
- Pielou’s evenness: *J′ = H′ ln S*

where *p*_*i*_ is the proportion of species *i* and *S* is the number of species.

The Uniform Manifold Approximation and Projection (UMAP) was applied to the relative abundance of bacterial species, capturing the first two dimensions with the R package umap (version 0.2.10.0).^77,78^ The two UMAP dimensions were then color-coded to represent two clinical groups (≤ 28 days on MV vs. deceased & > 28 days on MV), the Shannon diversity index, expression values of the five most abundant phyla (e.g. *Firmicutes*, *Tenericutes*, *Proteobacteria*, *Bacteroidetes*, and *Actinobacteria*), and ventilation days.

The significantly differential abundance of bacteria was detected through the combination of the Mann-Whitney *U* test^79,80^ and ZicoSeq (R package GUniFrac, version 1.8)^81,82^ in 4157 bacterial species between two clinical outcome groups, utilizing the CLR-transformed bacterial relative abundance matrix. the following parameters were applied in the ZicoSeq analysis: is.winsor=F, is.post.sample=F, feature.dat.type=“other”, link.func=list(function (x) x), perm.no=10000, while all other settings remained at their default values. Volcano plots of both results were created separately. The log_2_ fold-change used in the volcano plot for Mann-Whitney *U* test was calculated as follows: log_2_ e^Q2i (deceased & >28 days on MV) − Q2i (≤28 days on MV)^, where the median abundance (Q2_i_) was calculated for each bacterium (*i*) in the two clinical groups. Due to the log scale of the CLR transformation is based on the natural logarithm, necessitating an exponential back transformation followed by a log_2_ transformation of the values.

### Identification of the host DEGs and DEGs-associated bacteria

DEGs were detected from the filtered count matrix, which consisting of 16282 genes and 86 samples, using the R package edgeR (version 3.40.2)^64^ to determine the DEGs between the ≤ 28 days on MV group and the deceased & > 28 days on MV group. The criteria set as |log_2_ *FC*| ≥ 1 and FDR < 0.05. Spearman’s correlation coefficients (SCC)^83,84^ were applied between 16282 genes and 4157 bacterial species from 86 samples to determine the correlations between bacterial species (CLR-transformed bacterial relative abundance matrix) and host transcriptomic profiles (edgeR normalized log_2_-CPM matrix) by R package Hmisc (version 5.1-2).^85^ Fisher’s exact test was used to determine the bacteria that might be involved in perturbations of the host DEGs, focusing primarily on those that showed higher or lower associations with DEGs than with non-DEGs. The contingency table summarized whether there was a correlation with DEGs or with non-DEGs and whether the absolute value of the correlation was greater than 0.4 or not. The odds ratio (OR) and *p*-value were calculated as follows (using the R stats function “fisher.test”):

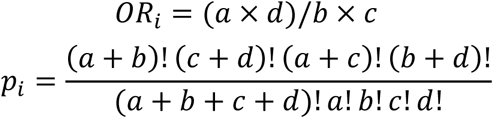

where *p_i_* is the *p*-value of bacteria *i*, *a* is the number of DEGs with |SCC| > 0.4, *b* is the number of non-DEGs with |SCC| > 0.4, *c* is the number of DEGs with |SCC| ≤ 0.4, and *d* is the number of non-DEGs with |SCC| ≤ 0.4. Multiple testing correction was applied to adjust *p*-values, and DEGs-associated bacteria were selected based on the criteria OR > 1 and FDR < 0.05.

### Discovery of bacteria relevant to patient survival from bacteria of differential abundance and bacteria associated with DEGs

Survival analysis was conducted using the R package survival (version 3.3-1).^86,87^ The log-rank test was employed to assess the impact of bacterial abundance on the survival probability in the subgroups (high vs. low) defined by the median CLR-transformed relative abundances of potentially and DEGs-associated bacteria or differentially abundant bacteria in the 86 BAL patients. Univariate analysis of survival was visualized using a Kaplan-Meier plot (survminer, version 0.4.9)^88^ when the *p*-value from the log-rank test results was less than 0.05. Continuous factors were converted into a binary classification (high (> median) vs. low (≤ median)) based on their median values.

### Inferred biological functions of the DEGs-associated, differentially abundant and survival-related bacteria

To elucidate the possible biological processes a bacterium is involved in, fast gene set enrichment analysis (FGSEA, fgsea version 1.24.0)^89^ was conducted using the rank of bacteria-correlated gene expression in descending order of the SCC values. The gene sets of Gene Ontology (GO) Biological Process from the Molecular Signatures Database (MSigDB) were imported and processed through the R package msigdbr (version 7.5.1).^90–93^ Due to the skewed distributions of SCC in the pre-ranked gene list, indicating a predominance of either positive or negative correlations, FGSEA was unable to generate a valid null distribution and enrichments for the gene sets on the smaller side of these severely unbalanced distributions. Consequently, the descending order of genes in the pre-ranked gene list was maintained, and randomly once-permuted SCC values in descending order between host genes and the CLR-transformed bacterial relative abundances were used as input for the pre-ranked gene list. The parameters used in the fgsea function were set as follows: minSize = 15, and maxSize = 500. The aggregation of selected GO terms based on the first twenty (twenty five), ten, or five absolute NES scores in descending order for each nine DEGs-associated (with two potentially DEGs-associated bacteria), the first ten up/down differentially abundant bacteria, and the 39 survival related bacteria was visualized in heatmap plots using the R package ComplexHeatmap (version 2.14.0).^68^

### Bacterial co-abundance network

FastSpar (version 0.0.10),^63,94,95^ a fast and parallelizable implementation of the SparCC algorithm with an unbiased *p*-value estimator, was used to calculate the correlation and significance between bacteria present (count > 0) in more than 50% of the BAL samples in each clinical outcome group on Ubuntu 18.04.6 LTS machine. Due to differences in sample size between the two clinical outcome groups (21 in the ≤ 28 days on MV group and 65 in the deceased & > 28 days on MV group), 21 samples from the group of deceased & > 28 days on MV were randomly selected to ensure comparability between the two co-abundance networks. Species pairs with an absolute value of SparCC correlations ≥ 0.5 and a *p*-value ≤ 0.001 in each clinical outcome group were isolated and visualized using Cytoscape (version 3.9.1).^19^ Molecular Complex Detection (MCODE, version 2.0.2), a plugin in Cytoscape, was employed to identify modules with tightly connected local regions using default settings^18^: degree cutoff = 2, node score cutoff = 0.2, K-core = 2, and max depth = 100. To ensure that the nodes were positioned identically in both networks, a subsequent merging process was applied to the two co-abundance networks. This process involved integrating the nodes and edges from both networks, which were then utilized as the foundational structure and visualized using Cytoscape. MCODE identified modules within each individual network that contained species of interest, which were marked with distinct colors and highlighted. The bacteria of interest were assigned color labels and different node shapes, including the top five abundant bacteria in the two clinical groups, the five most differentially abundant bacterial species enriched in each group, nine DEG-associated bacteria, and 39 survival-related bacteria. The highlighted clusters are represented by larger, colored nodes and thicker edges, while the remaining nodes are depicted as smaller light-gray nodes with thinner, partially transparent edges.

### Construction of a phylogenetic tree

The file SILVA_138.1_SSURef_NR99_tax_silva.fasta.gz, containing 16S-rRNA gene sequences, was acquired from the SILVA database.^62^ To construct a phylogenetic tree for the nine DEGs-associated bacteria, the fasta sequences of all relevant strains were initially gathered. However, with a total of 280 sequences, visualizing them in a phylogenetic tree proved challenging. Therefore, at most only the first ten matched sequences from each of the nine DEG-associated bacteria were selected for analysis. A total of 240 labeled bacteria were identified from two bacterial co-abundance networks, with first-hit fasta sequences extracted from 216 of these, sourced from 15,943 matched SILVA records. Given the impracticality of incorporating all sequences into the phylogenetic tree and the notion of similarity within the same genera, a random selection was conducted, choosing the first-hit fasta sequence for each genus from the 216 available records. In cases where the SILVA fasta database lacked records and no matching genus was found during the initial search for *Flammeovirga sp. L12M1*, *Friedmanniella sagamiharensis*, *Janthinobacterium sp. 1_2014MBL_MicDiv*, and *Virgibacillus sp. Bac332*, the first matched genus records were retrieved again from the SILVA fasta. Consequently, *Flammeovirga kamogawensis*, *Friedmanniella sp. D1C11*, *Janthinobacterium sp. MRL-AN3*, and *Virgibacillus marismortui* were used instead. A total of 95 bacterial fasta sequences were ultimately employed to construct the phylogenetic tree for the bacteria labeled within the co-abundance networks. The phylogenetic trees were efficiently generated and obtained through the one-click mode of Phylogeny.fr.^96^

### Drug prediction

To identify potential compounds for inhibiting COVID-19 based on the influence of various bacteria associated with COVID-19 survival and enrichment, we adopted a gene-expression-based approach to assess gene perturbation across multiple cell types under different compound treatments.^20^ In this study, we selected compounds by evaluating their efficacy using two strategies: (1) enhancing the expression of genes related to beneficial bacteria while preventing the downregulation of these genes; and (2) inhibiting the upregulation of genes associated with harmful bacteria while promoting the downregulation of these genes. The selection of these strategies was based on whether the input gene expression profile pertained to beneficial or harmful bacteria. In our context, beneficial bacteria were identified as those associated with a high survival probability subgroup in COVID-19 patients and enriched in those requiring MV for ≤ 28 days. Conversely, harmful bacteria were those associated with lower survival probabilities. Any bacterium did not fit the definition above was excluded from the further prediction. For drug prediction, we used data including log_2_ fold change, log_2_ CPM and FDR from bacteria-related DEGs for each bacterium (Table S10). The efficacy of compounds in the prediction process was quantified as a therapeutic score, with higher scores indicating more favorable compound performance. Two prediction models, d6 and d24, representing drug treatments for 6 hours and 24 hours, respectively, were employed. The prediction results were displayed using hierarchical clustering in a heatmap with the R package ComplexHeatmap (version 2.15.4).

## QUANTIFICATION AND STATISTICAL ANALYSIS

Utilizing the R base package stats, the two-sided Mann-Whitney *U* test was applied to compare the statistical differences of the CLR-transformed bacterial relative abundance between two clinical outcome groups.^97^ Given that the bacterial abundance matrix is typically sparse and does not conform to a normal distribution, the Mann-Whitney *U* test was deemed appropriate for this analysis. The log-rank test was employed to assess the impact of CLR-transformed bacterial relative abundance on the survival probabilities in high and low subgroups, and visualized with a KM plot.^87,88^ Differences were considered significant at a *p*-value of less than 0.05. The Benjamini−Hochberg (BH) method was used to correct *p*-values, addressing the multiple testing issue by calculating false discovery rates (FDRs) for DEGs-associated bacteria and drug prediction, with differences deemed statistically significant when the adjusted *p*-value fell below 0.05.

## Notes

### Competing Interest Statement

The authors have declared no competing interest.

