## Supplementary material for "Airway microbiome–host transcriptome networks link microbial dysbiosis to survival outcomes and therapeutic opportunities in severe COVID-19": Document S1.pdf

### SUPPLEMENTAL FIGURES

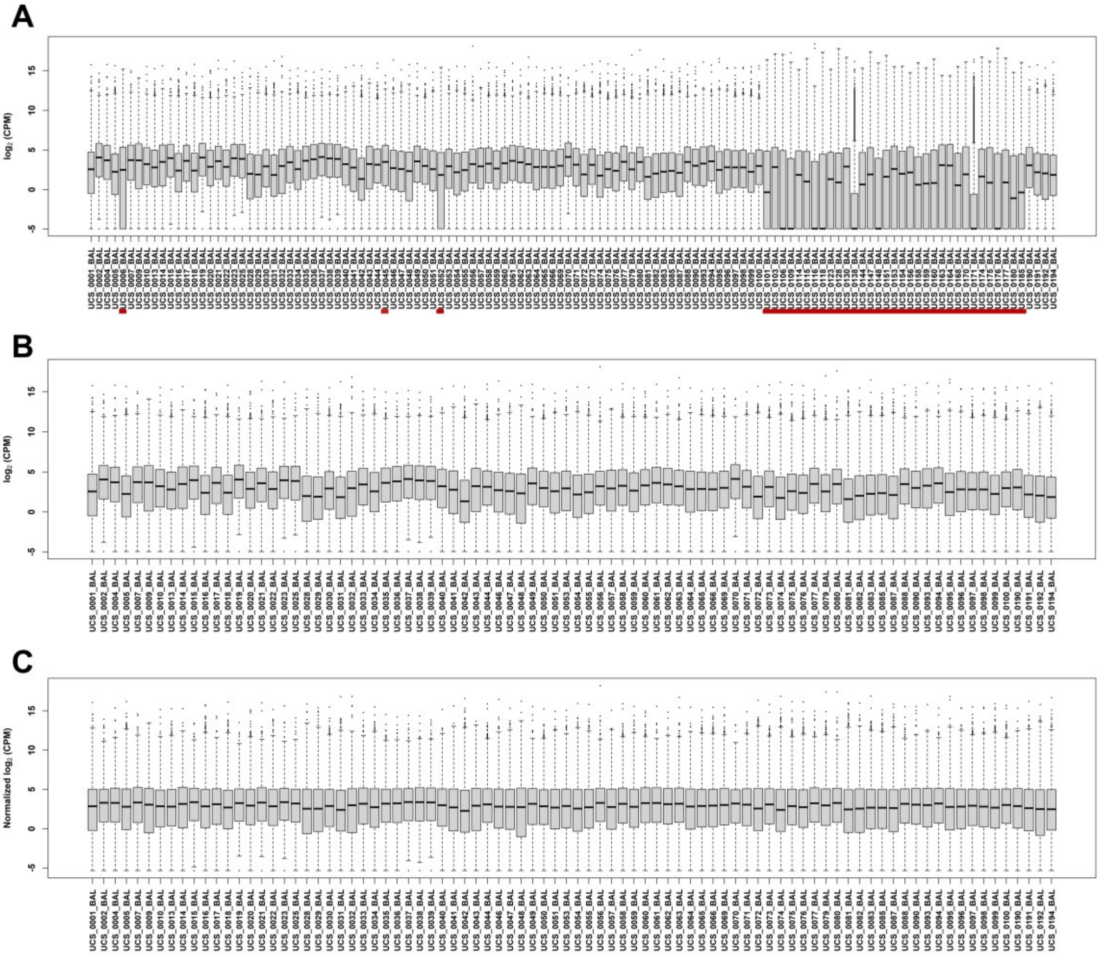


**Figure S1. Sample filtration based on the host transcriptome along with pre- and post-edgeR normalization.**

The host transcriptome underwent a few filtration processes to remove gene that were not lncRNAs or protein-coding genes, as well as expression levels that accounted for less than 20% of the 122 BAL samples. Boxplots were utilized to illustrate gene expression in all 122 BAL prior to (A) and following (B) the sample filtration process, along with subsequent edgeR normalization (C). The 36 samples that displayed a skewed gene distribution, low abundance, or lacked metadata information were highlighted in red (A) and excluded.

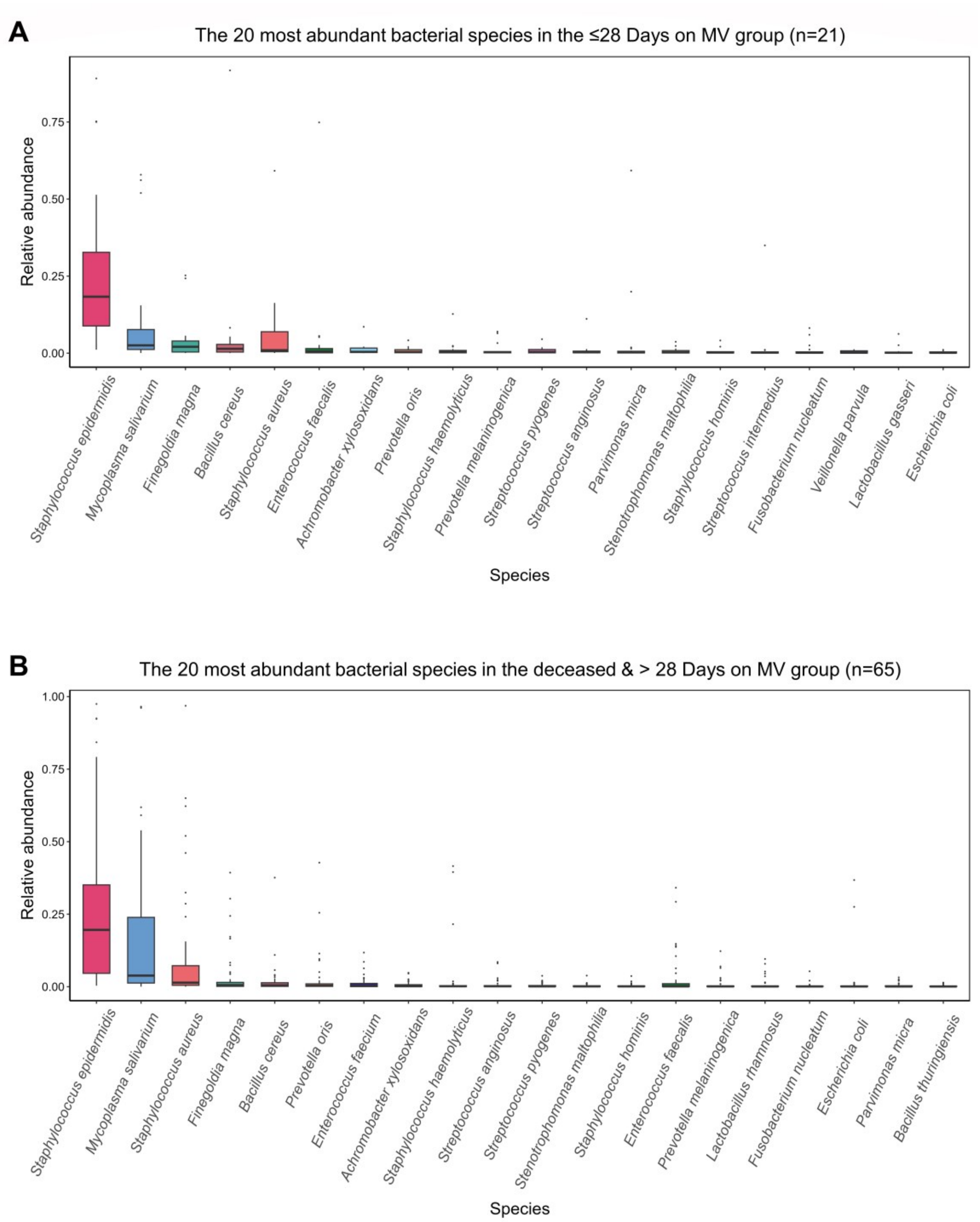


**Figure S2. The most abundant bacterial species, related to Figure 2.**

The top twenty abundant bacteria were identified in descending order of their median bacterial relative abundance in the two specified clinical outcome groups: (A) ≤28 Days on MV and (B) deceased & >28 Days on MV. Consistency was maintained by using the same colors for each respective bacterial species.

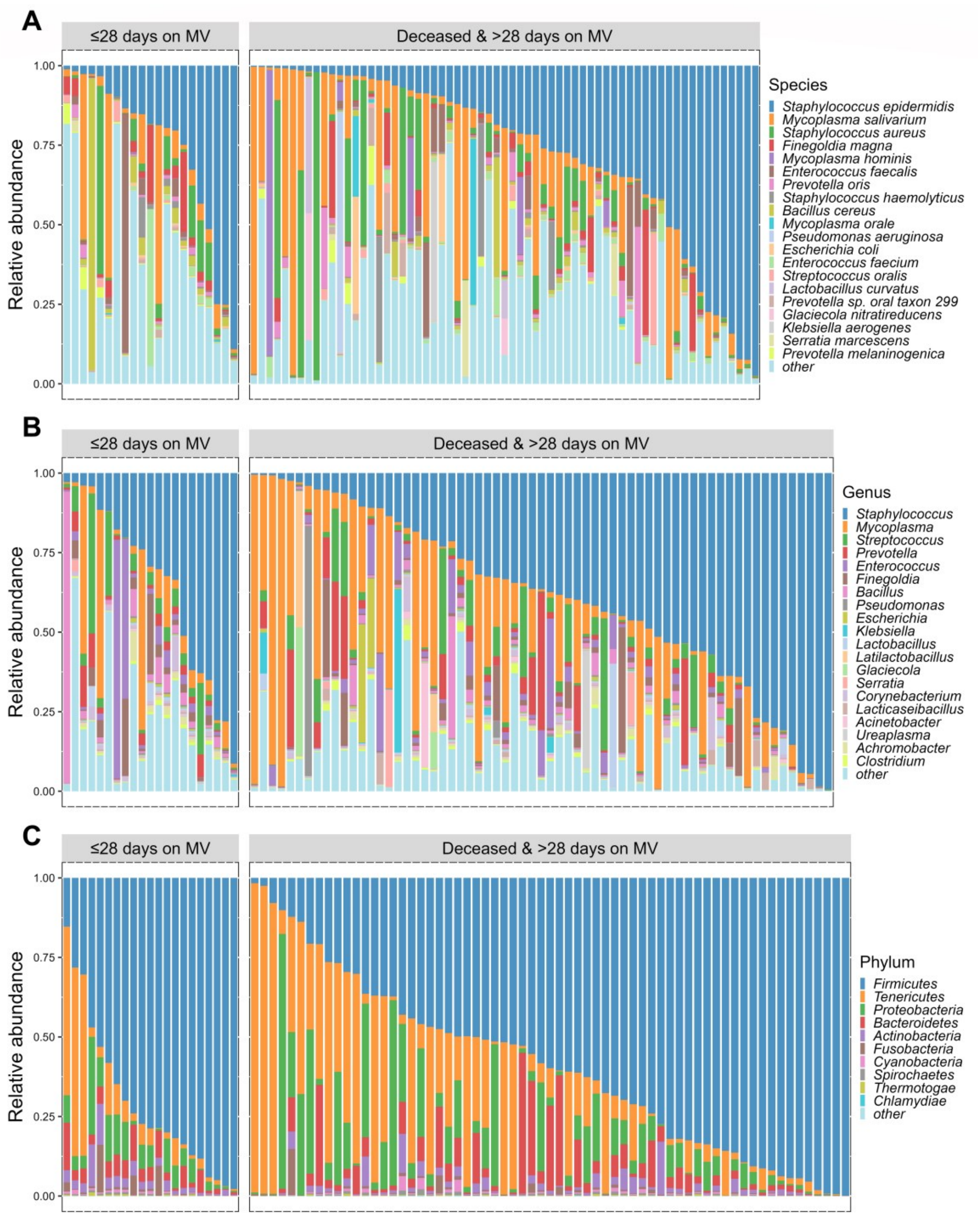


**Figure S3. The most abundant species, genera and phyla, related to Figure 2.**

The first twenty most abundant bacterial (A) species, (B) genera, and first ten most abundant (C) phyla, based on the average bacterial relative abundance within the deceased & >28 days on MV group, were identified and presented in the stacked barplots for two clinical outcome groups of all 86 BAL samples. The bacteria were arranged in descending order according to their mean relative abundance within the deceased & >28 days on MV group, and the columns representing the samples were ordered based on the most abundant bacteria at each taxonomic level.

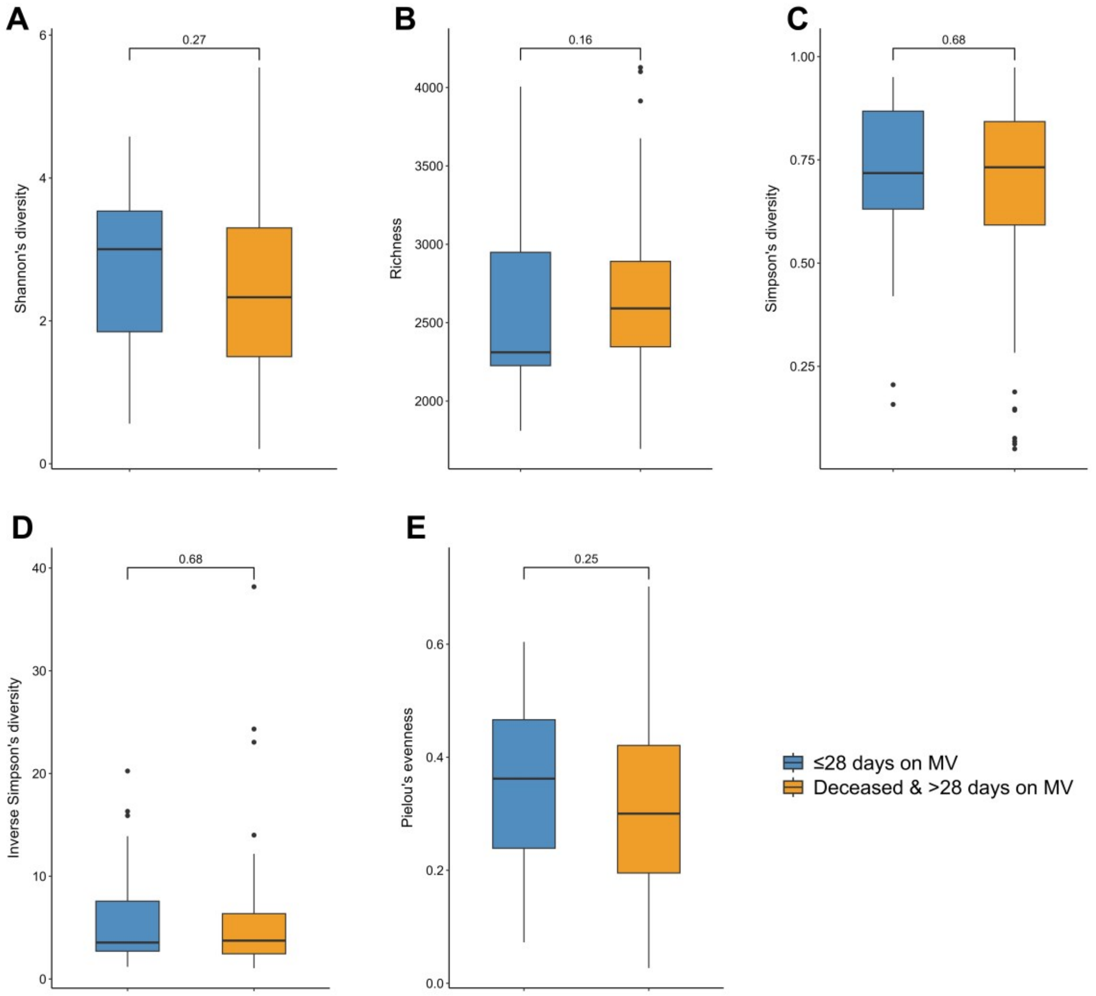


**Figure S4. The diversity indices and evenness, related to Figure 2.**

The group of deceased & >28 days on MV shows slightly reduction in diversity as indicated by the (A) Shannon index, (B) Richness, (C) Simpson and (D) Simpson reciprocal indices, as well as (E) Pielou’s evenness. However, the decrease does not reach statistical significance. The *p*-value was calculated using the Mann-Whitney *U* test.

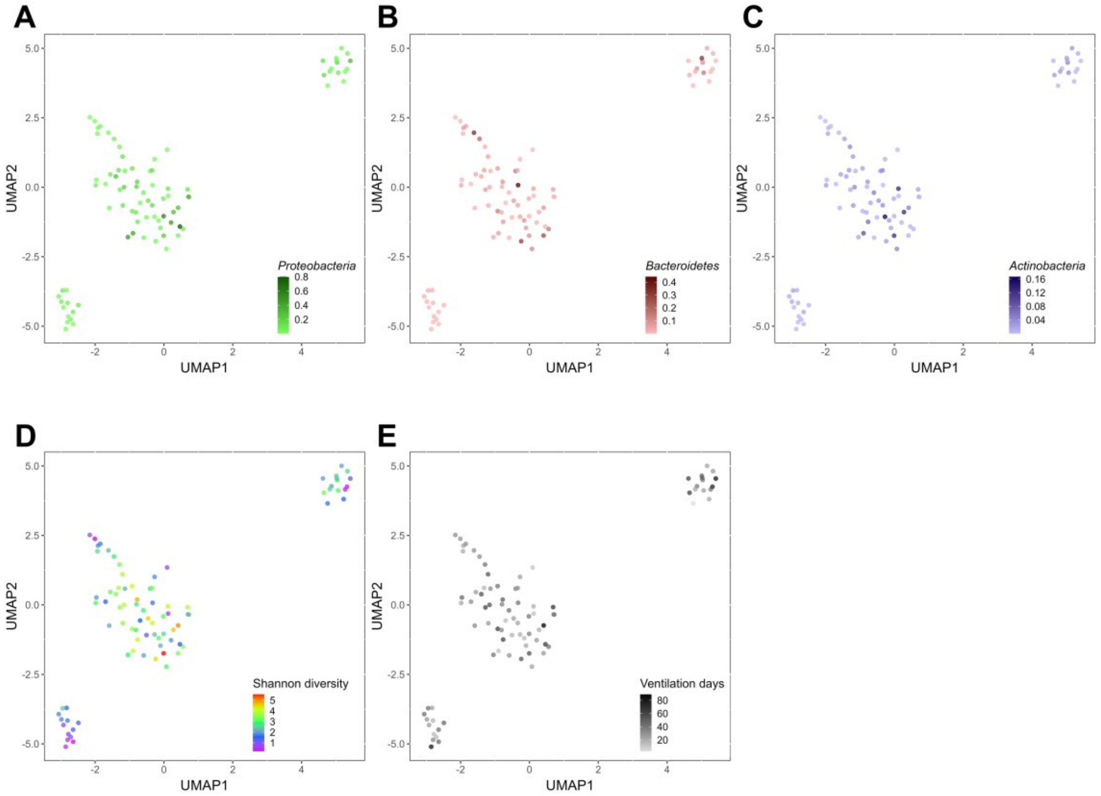


**Figure S5. The UMAP plots of 86 BAL samples, related to Figure 2.**

The UMAP plots presented herein were generated based on the relative abundance of bacterial species and were colored with the relative abundance of the third to fifth most abundant phyla (A) *Proteobacteria*, (B) *Bacteroidetes*, and (C) *Actinobacteria*, as well as (D) the Shannon index and (E) the duration of ventilation days.

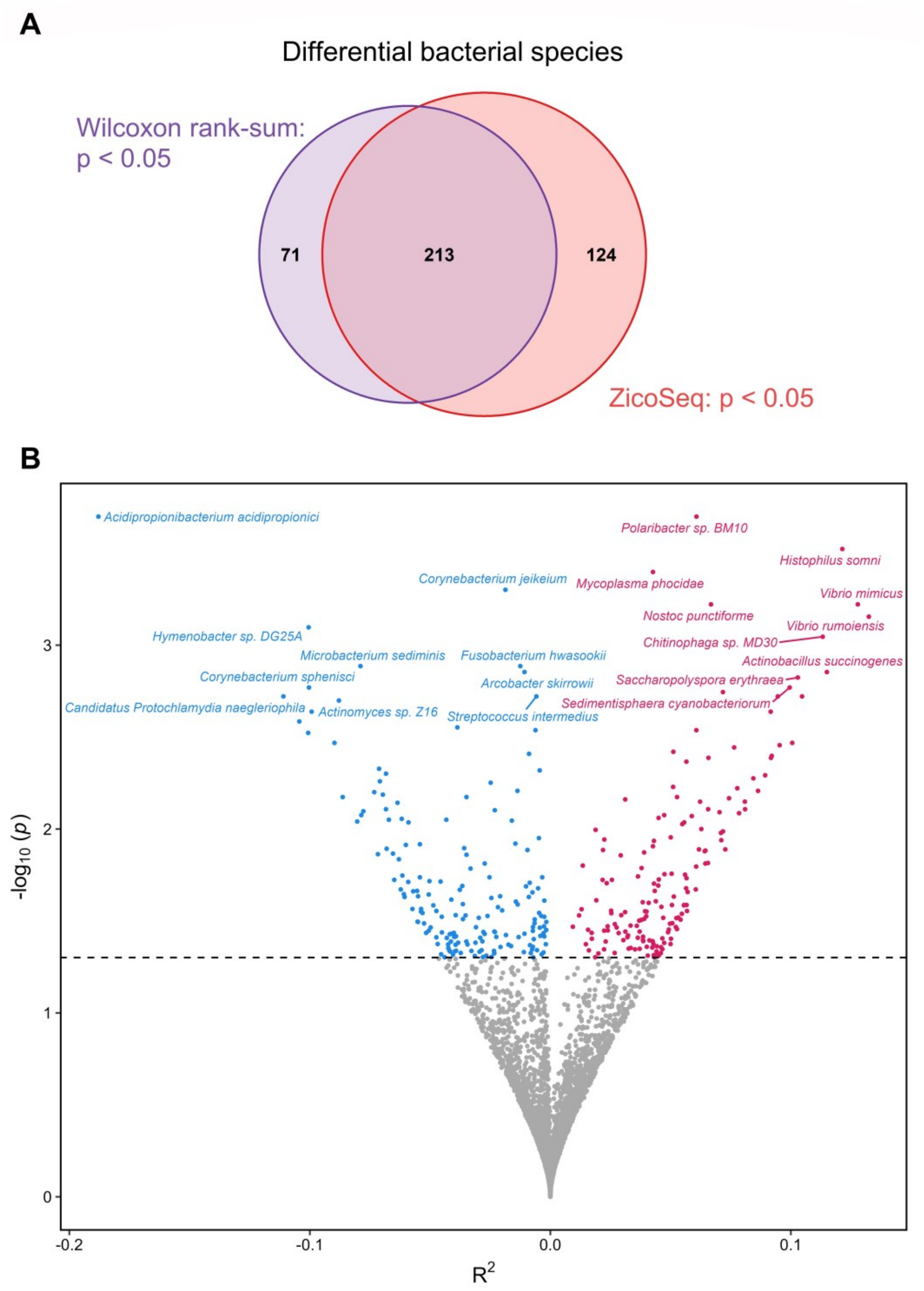


**Figure S6. Differential abundance of bacterial species, related to Figure 3.** (A) A total of 213 differential bacterial species were identified, as determined by the integrated findings of the Mann-Whitney test and ZicoSeq analysis, with both methods reaching a significance threshold of *p*-value < 0.05. (B) A total of 337 bacterial species demonstrated differential abundance, with 174 species exhibiting higher abundance in the ≤28 days on MV group (represented by dark sky blue color), whereas 163 of them were more enriched in the deceased & >28 days on MV group (indicated by dark pink color). The $R^2$ indicated the percentage of explained variance, while the *p*-value was calculated based on permutations, both of which were obtained from the ZicoSeq results.

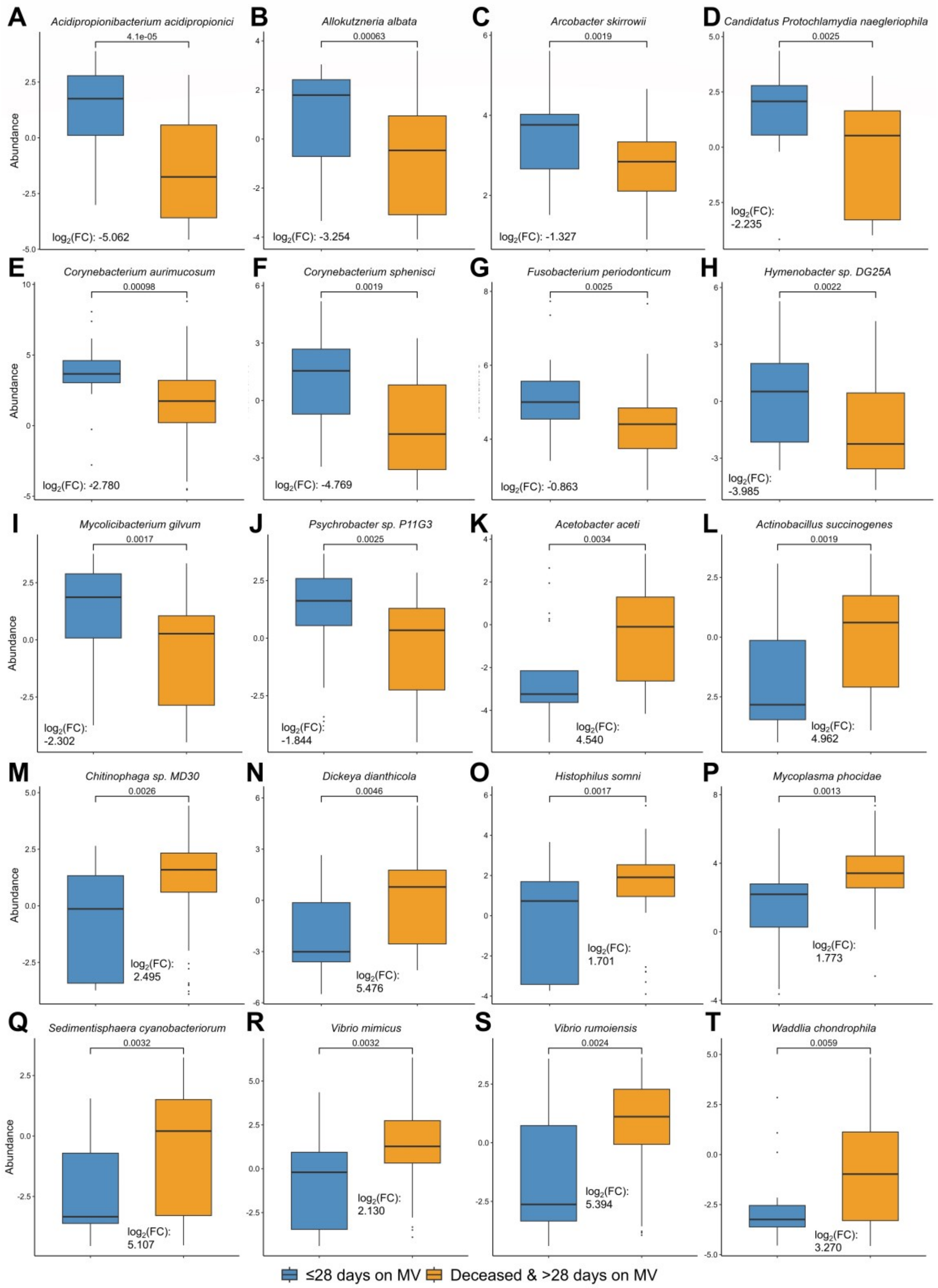


**Figure S7. The top ten significantly differential bacterial species enriched in each clinical group, related to Figure 3.**

(A–T) These twenty significantly and differentially abundant bacterial species were depicted with boxplots and they were arranged in alphabetical order of the species names in each group. The fold-change, represented as the ratio of exponential median difference between the CLR-transformed bacterial relative abundance between the two groups (deceased & >28 days on MV and ≤28 days on MV), was used to quantify the differences in abundance. The *p*-value was calculated using the Mann-Whitney *U* test.

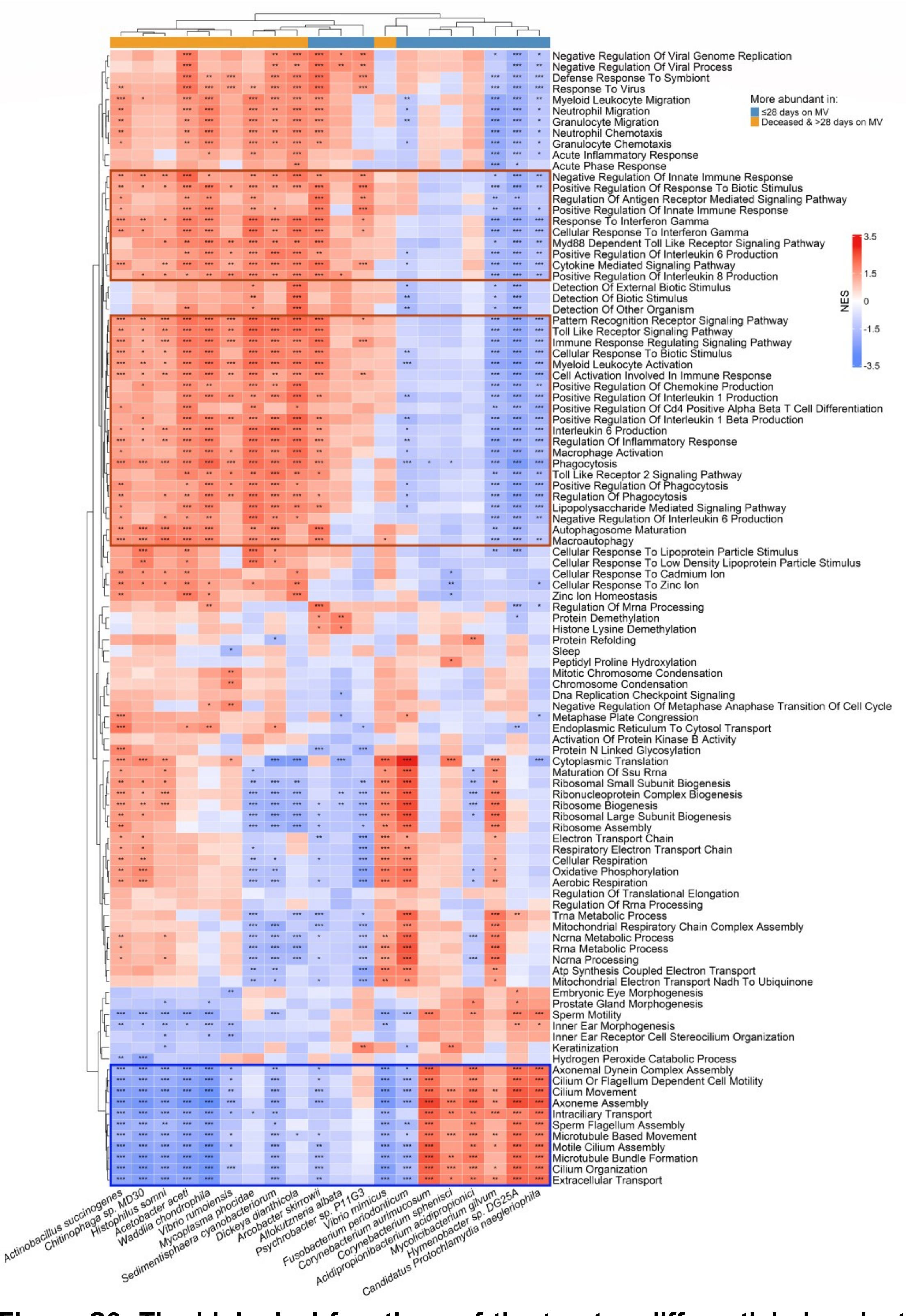


**Figure S8. The biological functions of the top ten differential abundant bacterial species enriched in two clinical groups, related to Figure 3.** The functions were derived from the union of the top ten absolute NES scores of GO terms across these twenty bacterial species and were presented in the form of the heatmap plot. The GO terms of each bacterium were labeled with an asterisk to indicate their significance as follows: *: 0.01 ≤ adjusted *p*-value < 0.05; **: 0.001 ≤ adjusted *p*-value < 0.01; ***: adjusted *p*-value < 0.001.

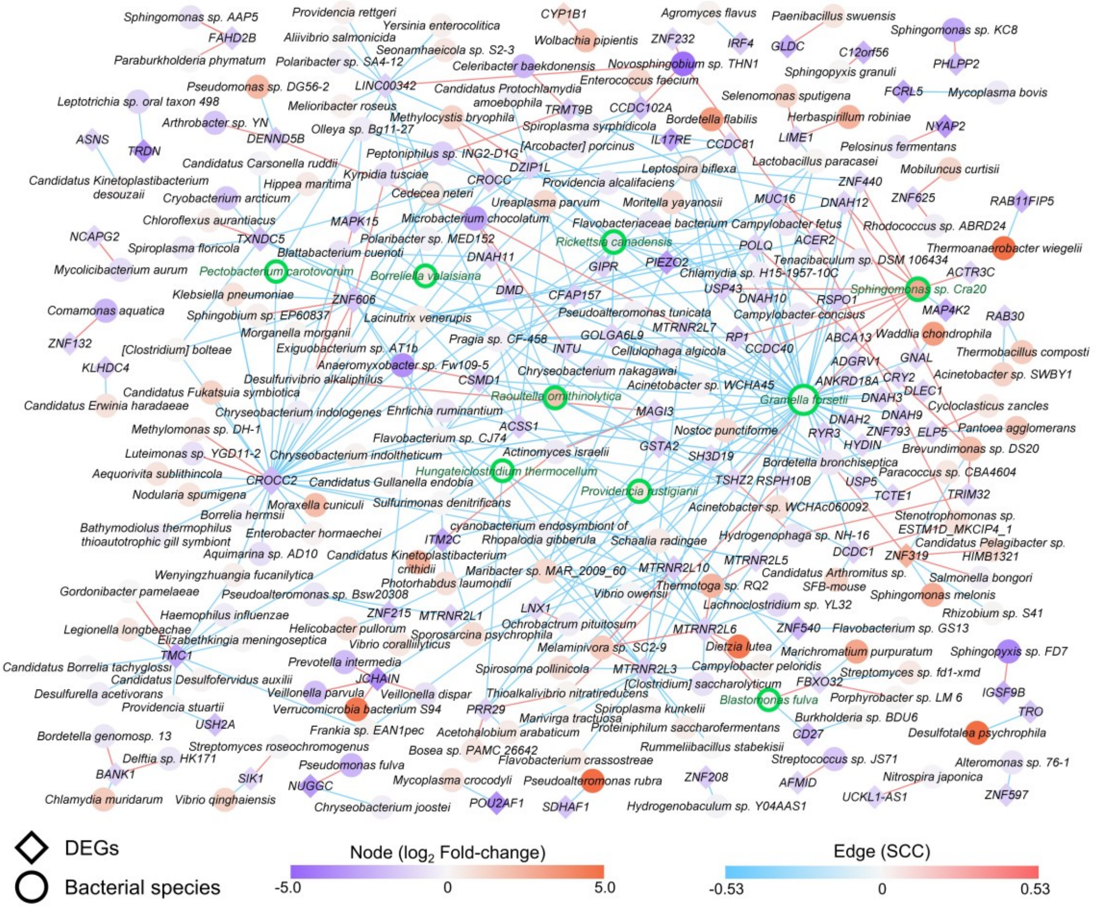


**Figure S9. The network of DEGs and their correlated bacterial species, related to Figure 4.**

DEGs and their correlated bacteria, with absolute SCC value greater than 0.4, were extracted and visualized as a network using Cytoscape. The DEGs were represented by diamond-shaped nodes, while the bacteria species were depicted as circular nodes. The fold-change of bacteria nodes is indicated by the ratio of exponential median difference between the two groups: deceased & >28 days on MV vs. ≤28 days on MV.

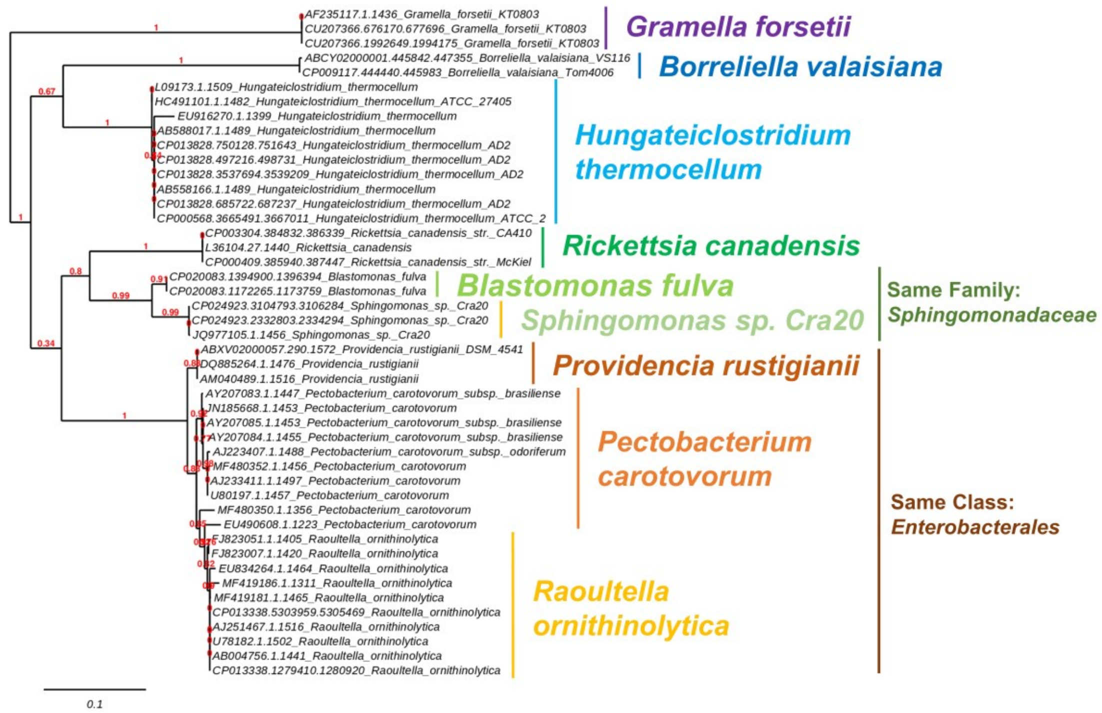


**Figure S10. The 16S-rRNA derived phylogenetic tree of the nine DEGs-associated bacterial species, related to Figure 4.**

The phylogenetic tree demonstrated the expected grouping of strains from the same species. Additionally, bacteria belonging to the same taxonomic category were also found to be clustered closely. For example, *B. fulva* and *S. sp. Cra20* were both categorized within the *Sphingomonadaceae* family. *P. carotovorum*, *P. rustigianii* and *R. ornithinolytica* were all classified under the *Enterobacterales* class.

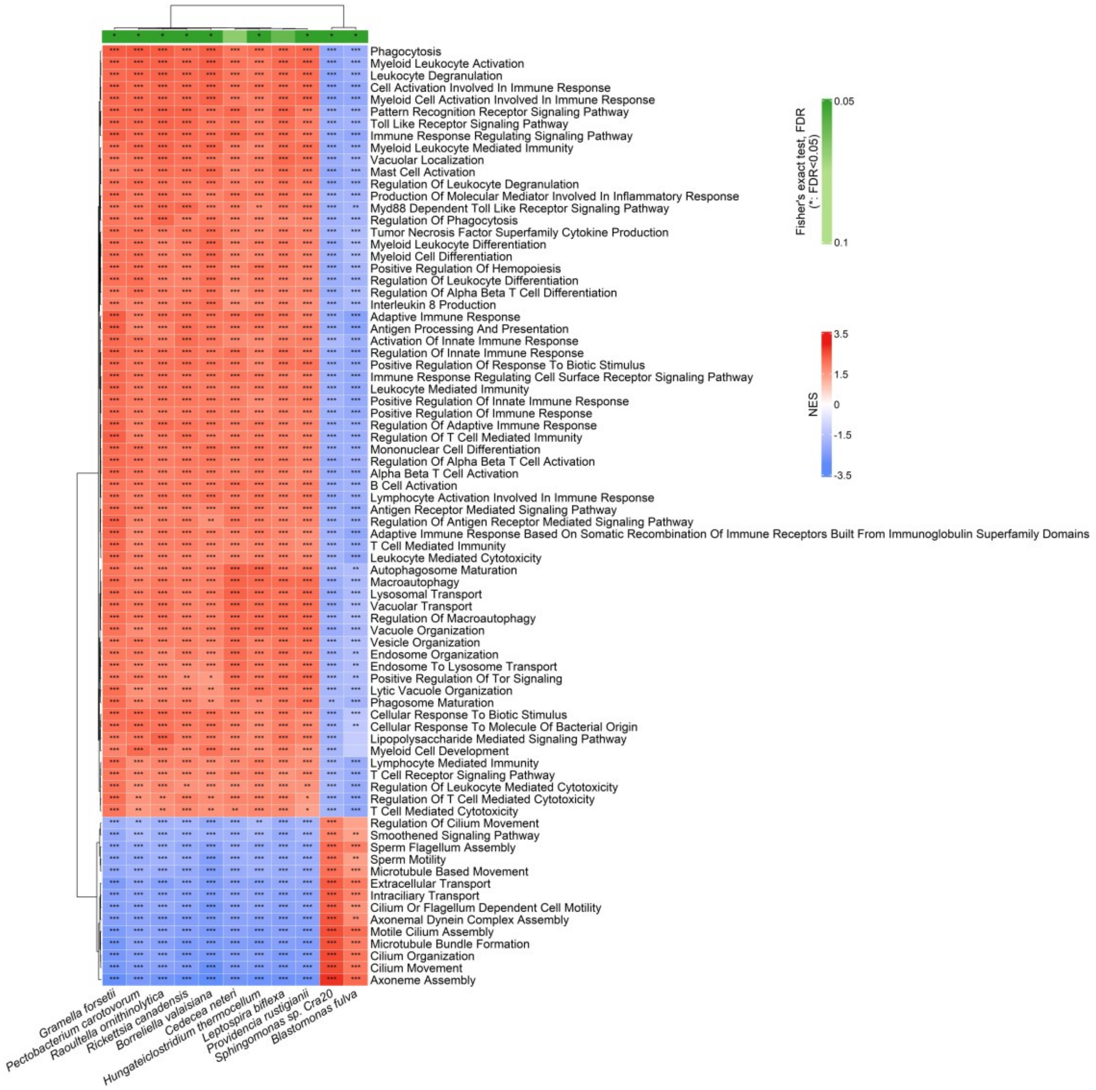


**Figure S11. The biological functions of the nine DEGs-associated bacteria and two potentially DEGs-associated bacteria, related to Figure 4.**

The functions were derived from the union of the top 25 absolute NES scores of GO terms across these eleven bacterial species and were presented in the form of the heatmap plot. The GO terms of each bacterium were labeled with an asterisk to indicate their significance as follows: *: 0.01 ≤ adjusted *p*-value < 0.05; **: 0.001 ≤ adjusted *p*-value < 0.01; ***: adjusted *p*-value < 0.001.

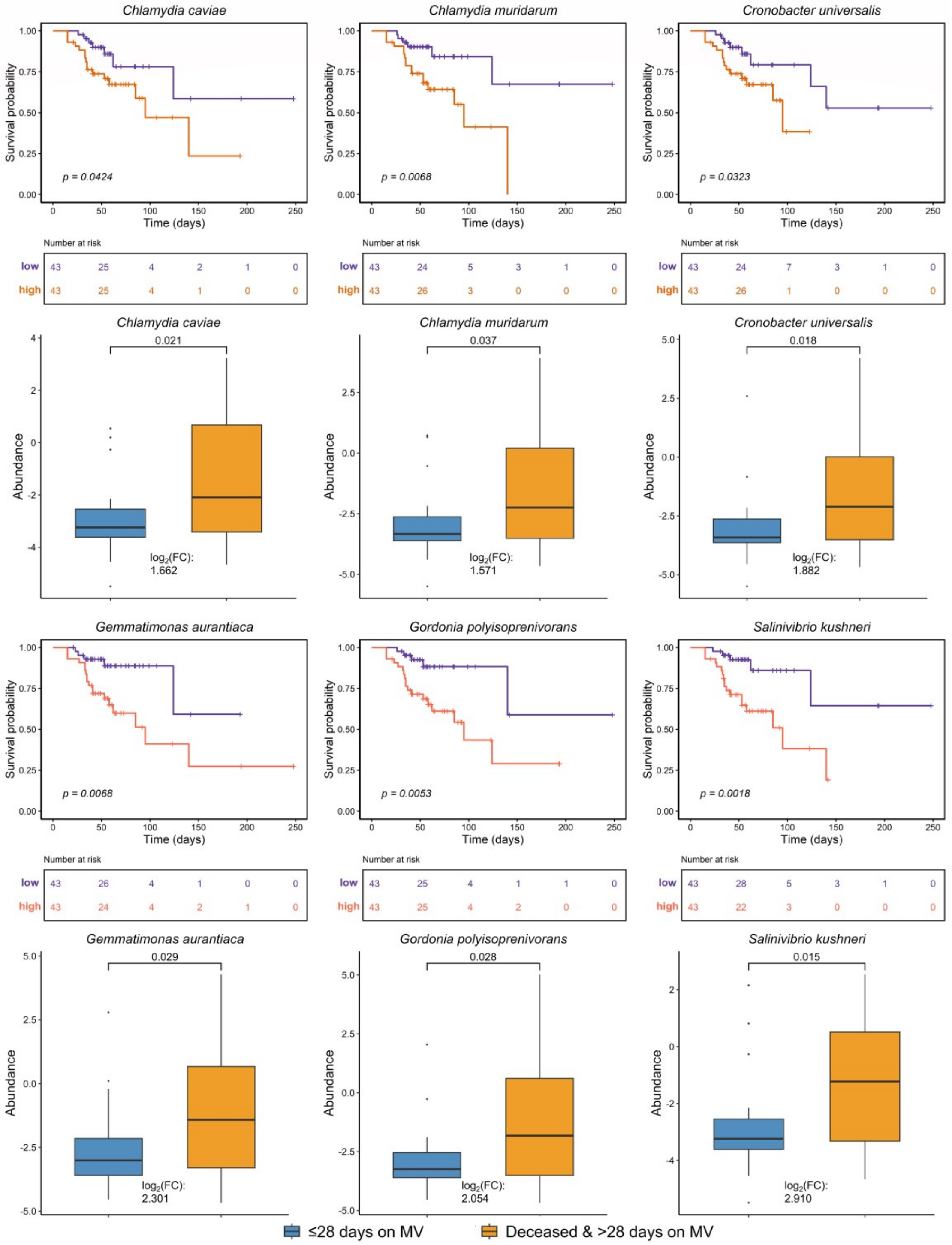


**Figure S12. Potentially hazardous types of bacteria in the severe COVID-19 patients, related to Figure 3 and 5.**

Severe COVID-19 cases exhibited a favorable survival outcome among patients belonging to the low abundant category of these microorganisms. Furthermore, these bacterial species were also found to be more abundant in the deceased & >28 days on MV group. The *p*-value in KM plot was computed by log-rank test. The fold-change, represented as the ratio of exponential median difference between the CLR-transformed bacterial relative abundance between the two groups (deceased & >28 days on MV and ≤28 days on MV), was used to quantify the differences in abundance. The *p*-value in boxplot was calculated using the Mann-Whitney *U* test.

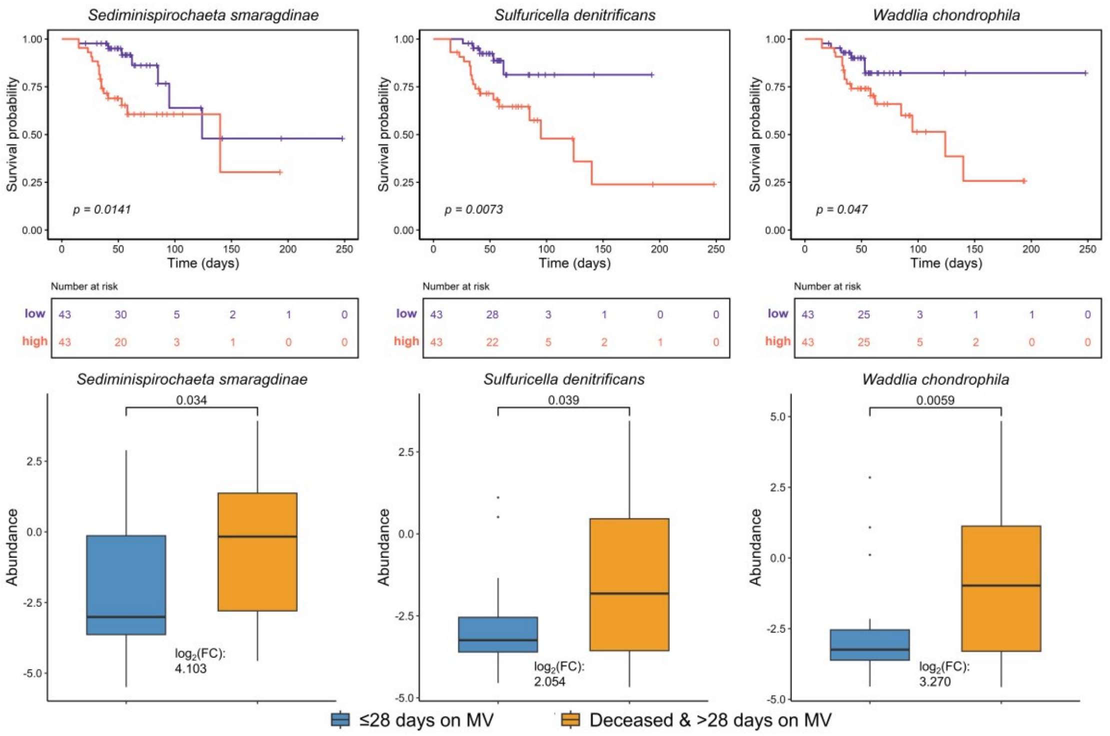


**Figure S12. Potentially hazardous types of bacteria in the severe COVID-19 patients, related to Figure 3 and 5. (Continued.)**

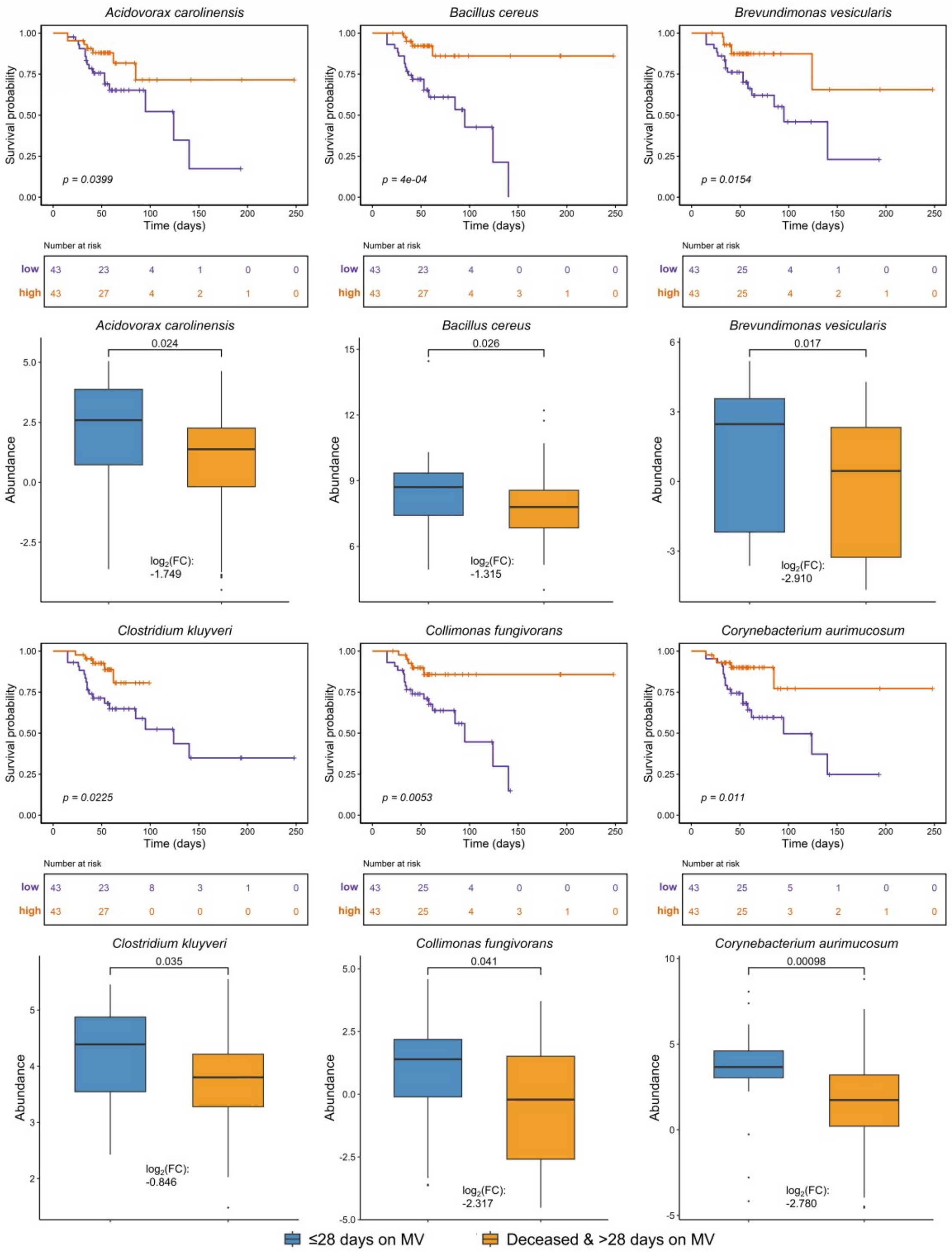


**Figure S13. Potentially advantageous bacterial species in the severe COVID-19 patients, related to Figure 3 and 5.**

Patients with a high abundance of these species exhibited better survival rates against those within a low abundance group of the bacteria. Furthermore, these bacterial species were also found to be more abundant in the ≤28 Days on MV group. The *p*-value in KM plot was computed by log-rank test. The fold-change, represented as the ratio of exponential median difference between the CLR-transformed bacterial relative abundance between the two groups (deceased & >28 days on MV and ≤28 days on MV), was used to quantify the differences in abundance. The *p*-value in boxplot was calculated using the Mann-Whitney *U* test.

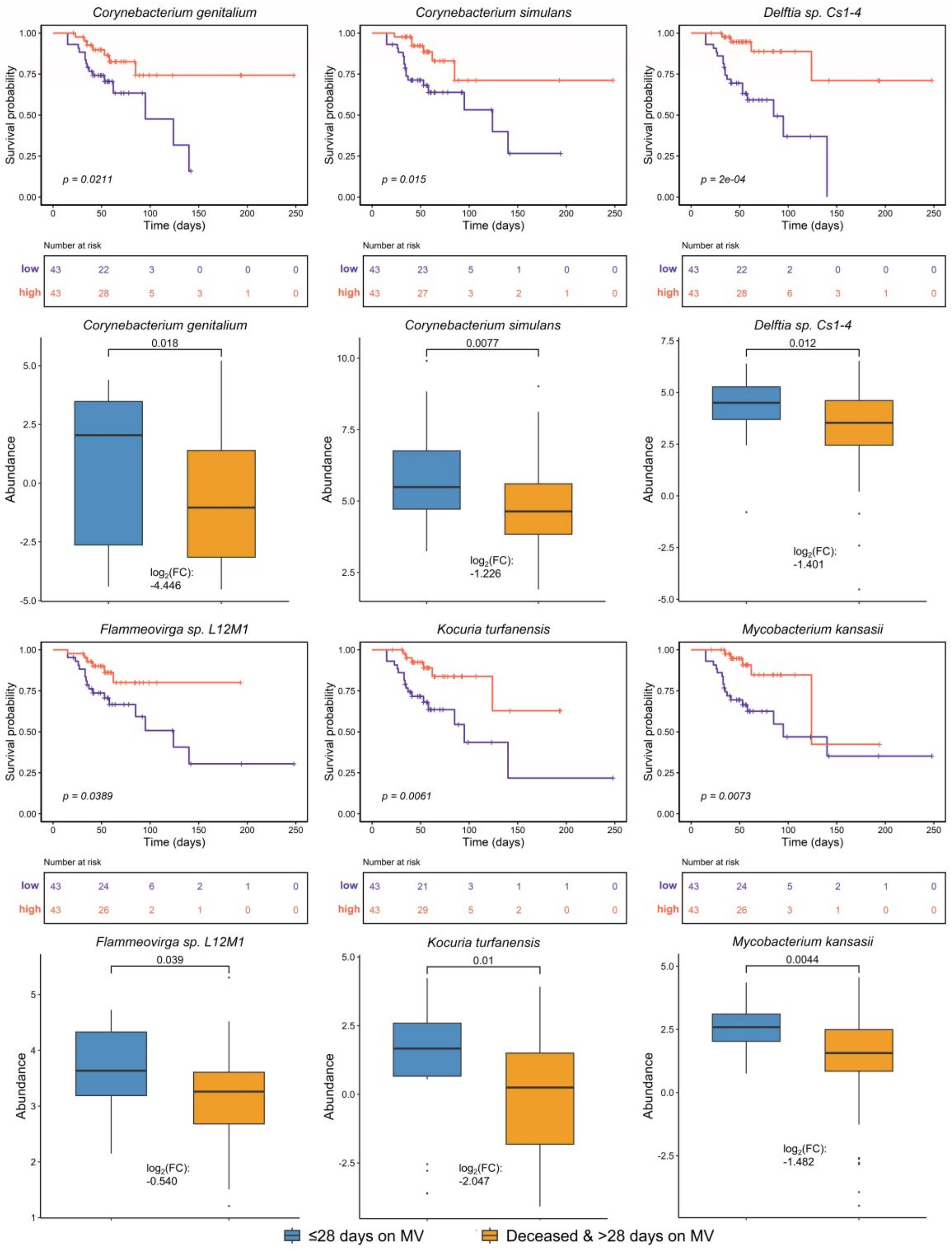


**Figure S13. Potentially advantageous bacterial species in the severe COVID-19 patients, related to Figure 3 and 5. (Continued.)**

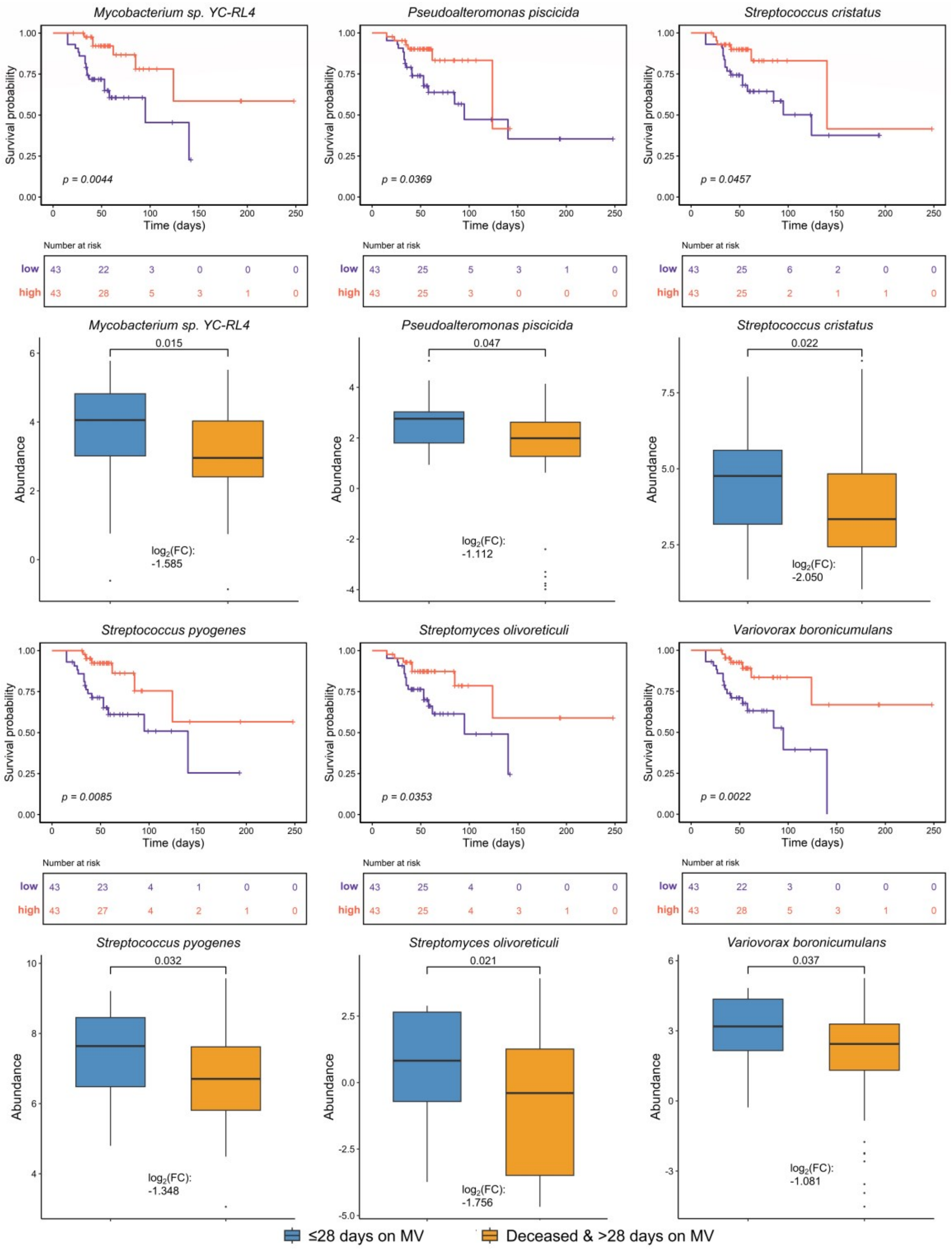


**Figure S13. Potentially advantageous bacterial species in the severe COVID-19 patients, related to Figure 3 and 5. (Continued.)**

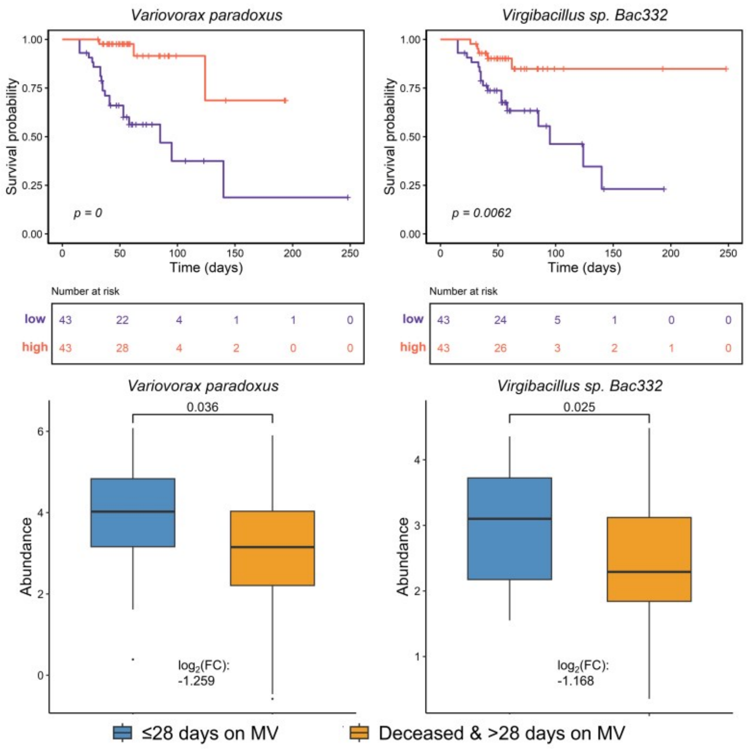


**Figure S13. Potentially advantageous bacterial species in the severe COVID-19 patients, related to Figure 3 and 5. (Continued.)**

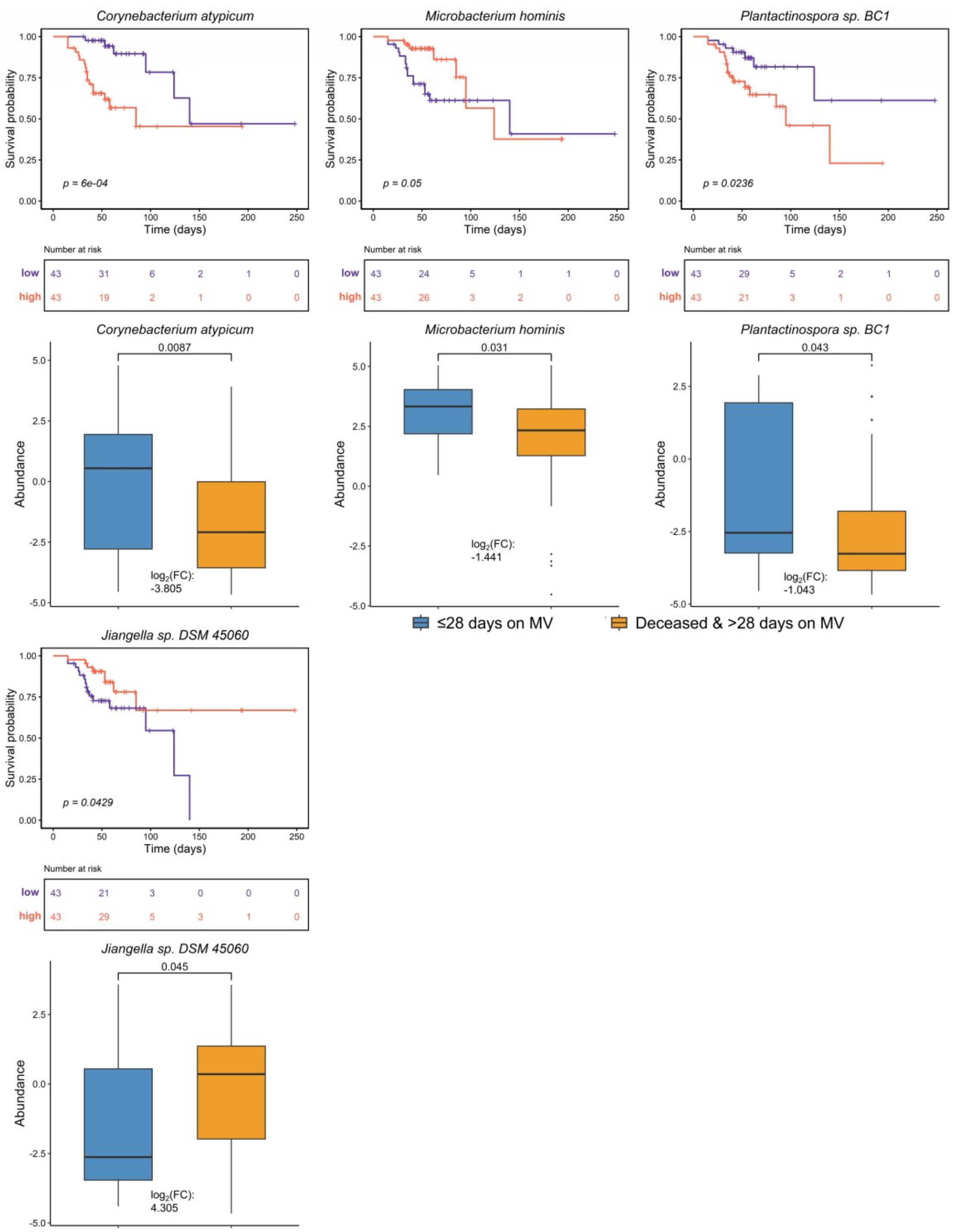


**Figure S14. The ambiguous types of bacteria in the severe COVID-19 patients, related to Figure 3 and 5.**

Patients with a greater abundance of *C. atypicum, M. hominis* and *P. sp. BC1* experienced a lower survival rate. Nevertheless, *C. atypicum, M. hominis* and *P. sp. BC1* exhibited higher abundance in the ≤28 Days on MV group. On the other hand, patients with a higher abundance of *J. sp. DSM 45060* displayed an improved survival rate, yet this bacterium was discovered to be more abundant in the deceased & >28 days on MV group. The *p*-value in KM plot was computed by log-rank test. The fold-change, represented as the ratio of exponential median difference between the CLR-transformed bacterial relative abundance between the two groups (deceased & >28 days on MV and ≤28 days on MV), was used to quantify the differences in abundance. The *p*-value in boxplot was calculated using the Mann-Whitney *U* test.

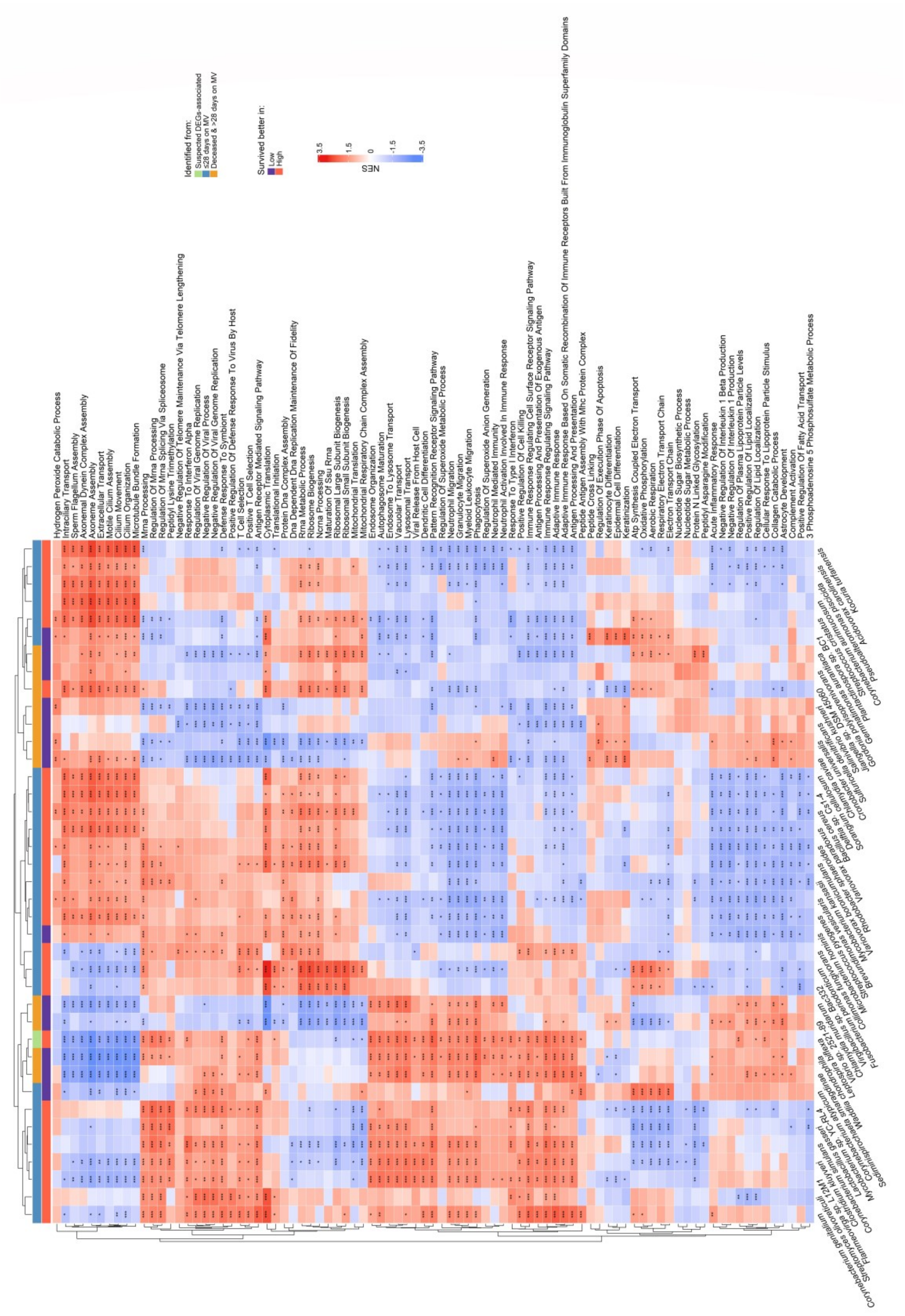


**Figure S15. The biological functions of the survival related bacterial species, related to Figure 4 and 5.**

The functions were derived from the union of the top five absolute NES scores of GO terms across the 39 survival-related bacterial species and were presented in the form of the heatmap plot. The GO terms of each bacterium were labeled with an asterisk to indicate their significance as follows: *: 0.01 ≤ adjusted *p*-value < 0.05; **: 0.001 ≤ adjusted *p*-value < 0.01; ***: adjusted *p*-value < 0.001.

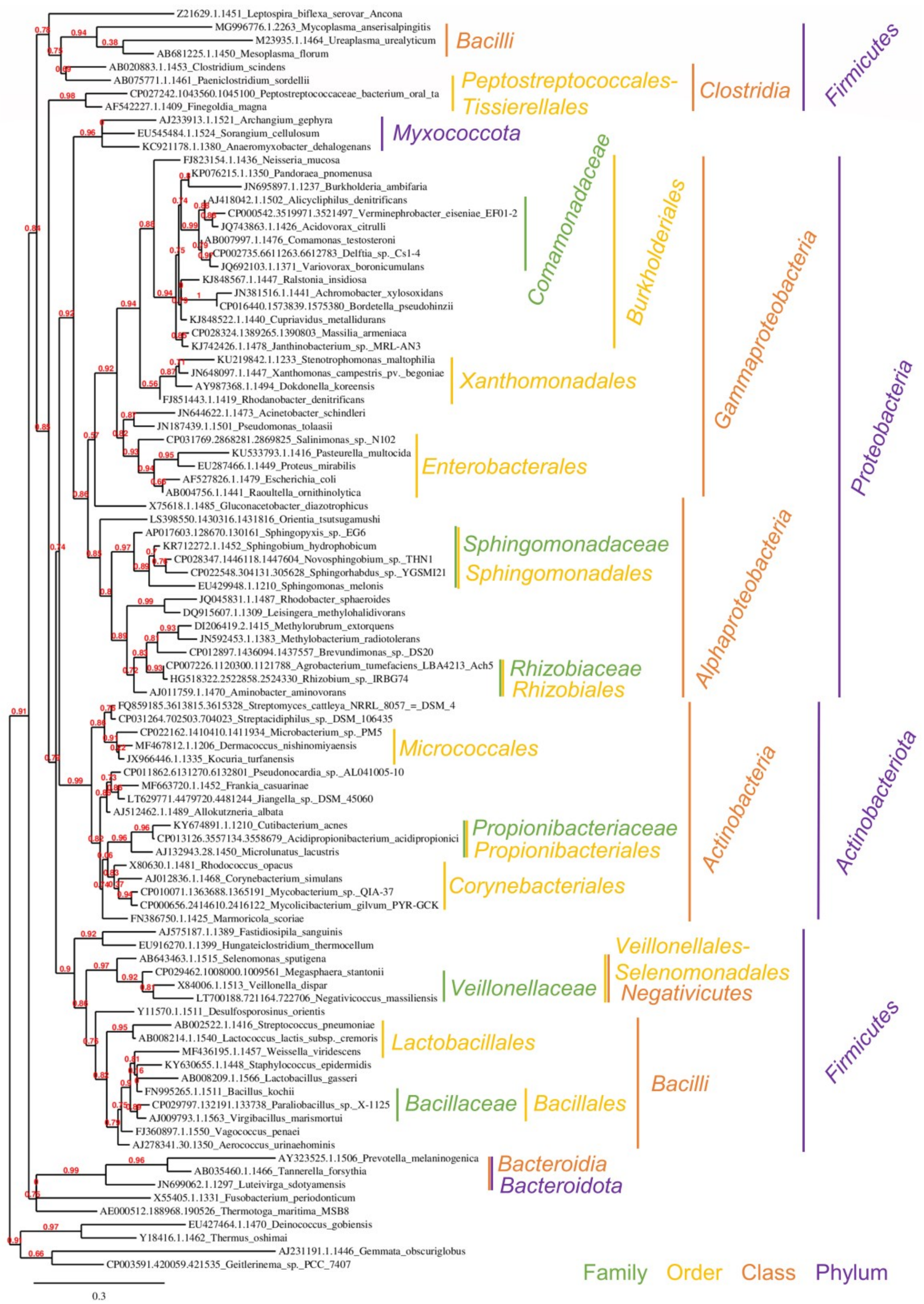


**Figure S16. The 16S-rRNA derived phylogenetic tree of nodes (bacterial species) in the highlight co-abundance networks, related to Figure 6.** Bacteria belonging to the same taxonomic category were clustered closely.

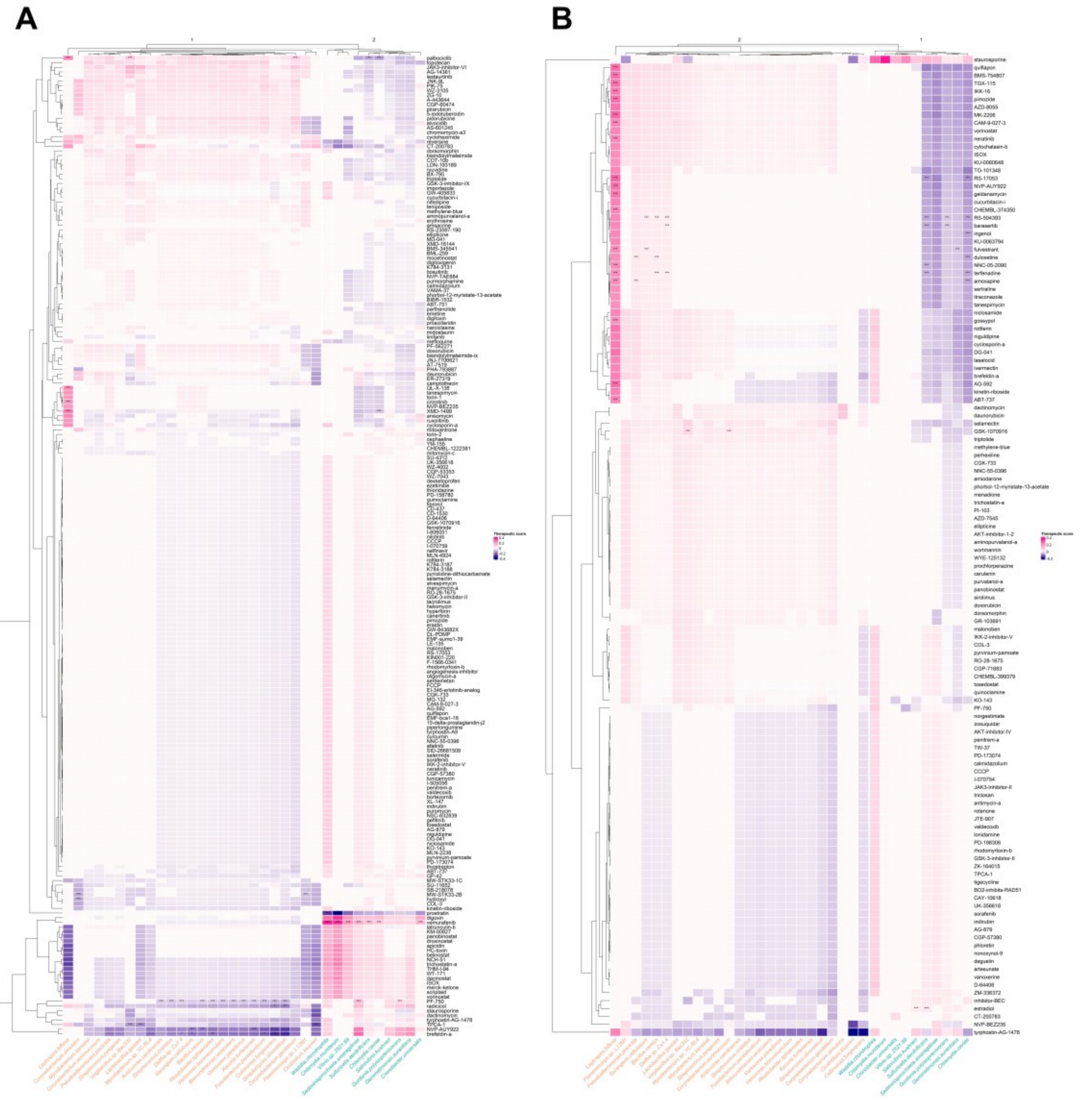

**Figure S17. Therapeutic score of complete compounds in the gene-expression-based drug prediction, related to Figure 7.**

Hierarchical clustering reflects the similarity of ability for gene perturbation between compounds and the relevance of bacterium-related gene expression between bacteria in (A) drugs for 6 hours (d6) prediction model and (B) drugs for 24 hours (d24) prediction model. The rows represent compounds in the prediction. The columns represent different bacteria-related expression profiles. Beneficial bacterium is labeled in orange; harmful bacterium is labeled in blue-green. The significant of the prediction is marked with *: 0.01 ≤ *p*-value < 0.05; **: 0.001 ≤ *p*-value < 0.01; ***: *p*-value < 0.001. Refer to Figure S17 for the therapeutic score of complete compounds.
